# Replication stress increases de novo CNVs across the malaria parasite genome

**DOI:** 10.1101/2024.12.19.629492

**Authors:** Noah Brown, Aleksander Luniewski, Xuanxuan Yu, Michelle Warthan, Shiwei Liu, Julia Zulawinska, Syed Ahmad, Nadia Prasad, Molly Congdon, Webster Santos, Feifei Xiao, Jennifer L Guler

## Abstract

Changes in the copy number of large genomic regions, termed copy number variations (CNVs), contribute to important phenotypes. CNVs are readily identified using conventional approaches when present in a large fraction of the cell population. However, CNVs in only a few genomes are often overlooked but important; if beneficial, a de novo CNV that arises in a single genome can expand during selection to create a population of cells with novel characteristics. While single cell methods for studying de novo CNVs are increasing, we continue to lack information about CNV dynamics in rapidly evolving microbial populations. Here, we investigated de novo CNVs in the genome of the *Plasmodium* parasite that causes human malaria. The highly AT-rich *P. falciparum* genome readily accumulates CNVs that facilitate rapid adaptation. We employed low-input genomics and specialized computational tools to evaluate the impact of sub-lethal stress on the de novo CNV rate. We observed a significant increase in genome-wide de novo CNVs following treatment with an antimalarial compound that inhibits replication. De novo CNVs encompassed genes from various cellular pathways participating in human infection. This snapshot of CNV dynamics emphasizes the connection between replication stress, DNA repair, and CNV generation in this important microbial pathogen.

## INTRODUCTION

Changes in the copy number of large genomic regions, termed copy number variations (CNVs), are a source of phenotypic diversity for many organisms (as reviewed in (1–3)), including rapidly evolving microbes such as bacteria, yeast, and viruses (4–6). CNVs also contribute to many diseases from cancer to blood, metabolic, neurological, and infectious diseases (7–19). Increased access to genome sequencing has facilitated the identification of these important genomic rearrangements, especially following selection. CNVs that are identified using standard “bulk” analysis approaches (e.g. read coverage methods) are present in a large fraction of the cell population (>50% (2)). However, those in a minority of genomes, or even a few genomes, are “averaged” away during analysis steps. This artifact limits our ability to assess how CNVs arise and contribute to the genomic diversity of individual cells within a population. In order to observe a genome’s evolutionary potential in the absence of selection, we require approaches specifically designed to detect CNVs that are not present in a predecessor “parental” genome. Due to their rare and novel nature, these events are termed “de novo” CNVs (3,20–24). Early experimental progress detecting de novo CNVs involved cloning individual cells, which takes time, is prone to contamination, and prevents detection of detrimental CNVs (20–22). Misalignment of short-reads to reference genomes (i.e. discordant or split reads) can also indicate the presence of de novo CNVs, but false positives are common if matched normal samples are not available (reviewed in (25)). Recent single cell-based methods have successfully assessed de novo CNVs during experimental evolution, disease progression, and tissue development (as reviewed in (25–29)). While the reach of single cell analysis is expanding, we continue to lack information about de novo CNVs in microbes and their dynamics in evolving pathogen populations. The *Plasmodium* parasite that causes malaria readily accumulates kb-sized CNVs in its genome (30–33). This protozoan propagates within the mosquito vector and then the human host, where it infects the liver and cyclically invades red blood cells (erythrocytic cycle, **Fig. 1A**). CNVs impact parasite survival, allowing evasion of clinical detection (34), expansion of beneficial gene families (35), invasion of new host cells (36), and development of antimalarial resistance (37–39). We are specifically interested in the genetic diversity of one species of malaria, *P. falciparum*, since this may explain its rapid adaptation to new drugs and host environments (40). Due to its relatively small, AT-rich genome (23 Mb, 19.4% GC (41)), low-input genomics is challenging in this single cell protozoan. However, we previously optimized a single cell genomics approach for erythrocytic stage *P. falciparum* parasites and recently developed novel computational tools to evaluate de novo CNVs in challenging genomes (42–44).

**Figure 1.**
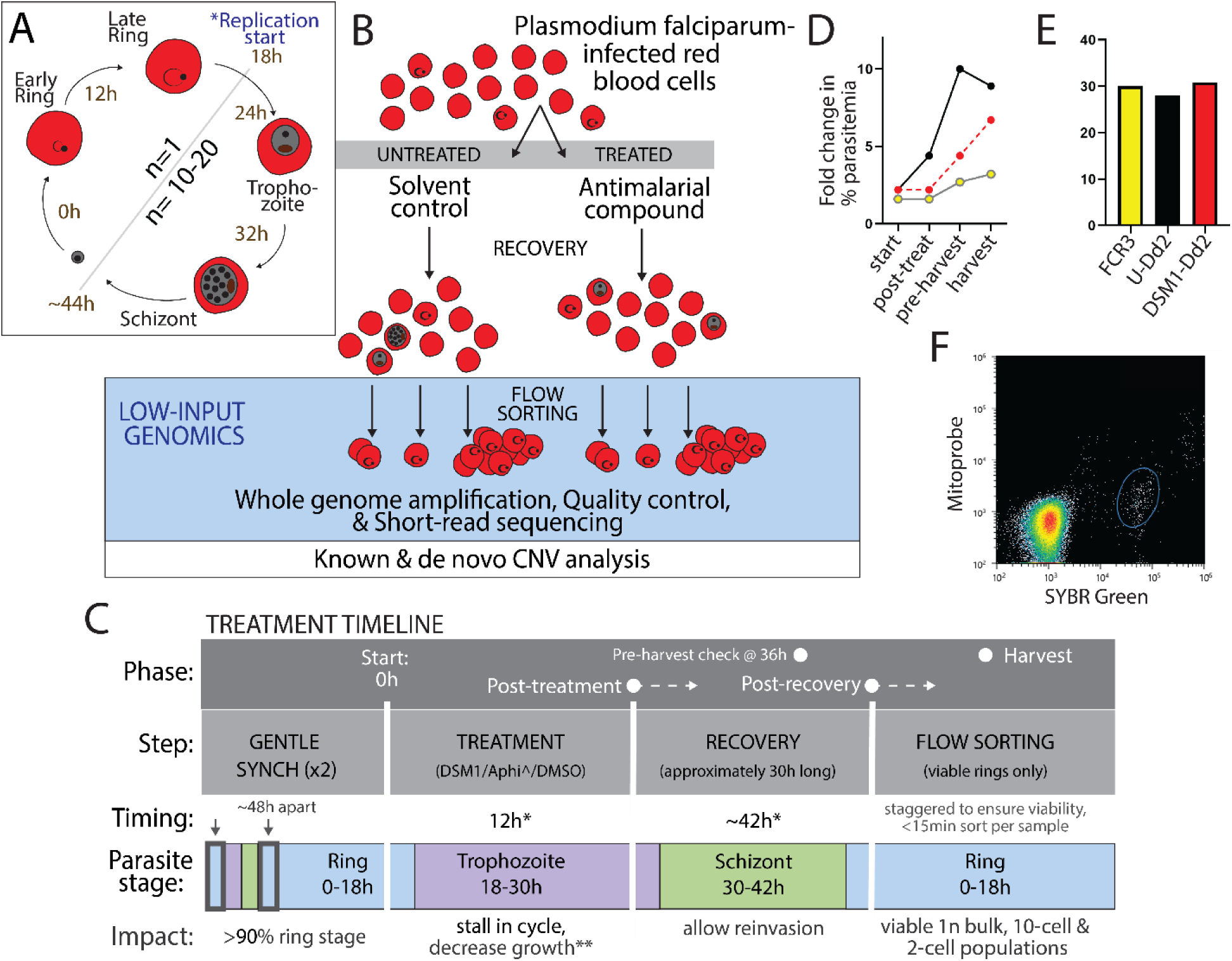
**Low-input genomics approach for analysis of malaria parasite genomes under stress**. **A**. *P. falciparum* erythrocytic cycle including approximate timing (h) and genome number (n). Cycle length is 44h total for the *Dd2* parasite line (78,79). Ring stage parasites have a single genome, replication begins at trophozoite stage, and schizonts develop with 10-20 individual parasites within a single erythrocyte. **B**. Overview of low-input genomics procedures. Early erythrocytic stage *P. falciparum* parasites grown in vitro were treated with an antimalarial compound (DSM1) or the solvent control (dimethylsulfoxide, DMSO) followed by recovery and reinvasion to produce a new round of ring stage parasites (see details in panel C). Parasite-infected erythrocytes were isolated using flow sorting (10-cell control or 2-cell samples) and genomes were amplified using a modified MALBAC-based whole genome amplification approach. Quality control steps involved DNA quantification to assess amplification success, droplet digital PCR to assess parasite genome amplification, and PCR-high resolution melting to assess sample cross-contamination. After Illumina short-read sequencing, reads were filtered, trimmed and used as input for copy number variation analysis using two approaches, HapCNV and LUMPY. **C**. Timeline of parasite treatment and recovery. For all experiments (pilot and low-input genomics), parasites were synchronized to enrich for rings before short-term treatment was applied. Parasite growth and viability were tracked from start, to post-treatment, to post-recovery, and harvest. The recovery period allowed for completion of the erythrocytic cycle and reinvasion to produce rings before harvest for low-input genomics (see more details on pilot treatments in **Fig. S1A** and low-input genomics in **Fig. S2A**). *, cumulative time from start (0h). **treatment led to a decrease in parasitemia and delay in ring progression and therefore staging is approximate. Aphi, aphidicolin. **D**. Mean growth rate of parasite lines and treatments for low-input genomics experiment: *FCR3* (yellow circles), untreated *Dd2* (black circles), and treated *Dd2* parasites (red circles). Fold change in % parasitemia was calculated by dividing the ending parasitemia from the starting parasitemia for the period being measured. Cumulative timing is as follows: start (0h), post treat (12h), pre-harvest (38h), harvest (+40-43h). N=2. Cumulative parasitemia and ring percentage shown in **Fig. S2B**; raw flow cytometry plots tracking viability and parasitemia shown in **Fig. S2C-G. E.** Recovery time for parasite lines and treatments: *FCR3* (yellow), untreated *Dd2* (black), and DSM1-treated *Dd2* parasites (red). Recovery is defined as the period just after treatment until harvest for down-stream processing. **F.** FACS plots at parasite isolation step for DSM1-treated *Dd2* samples. MitoProbe DiIC1 (5) (stains viable parasites) vs SYBR Green (stains parasite genome) indicates the proportion of erythrocytes (heat map of density) that contain viable ring stage parasites (blue circle: 0.38% ring stage parasitemia). Plots for other parasite lines are depicted in **Fig. S4**.

Here, we combined these advancements and made additional improvements to directly investigate de novo CNV formation in the *P. falciparum* genome. Since various types of cellular stress can induce genetic change (reviewed in (2,45–47)), we also evaluated the impacts of sub-lethal stress on de novo CNV development. Using a low-input genomics pipeline, we observed that replication stress increased the number of de novo CNVs across the parasite genome. This finding provides key evidence for adaptive amplification in an important pathogen that has evolved strategies to encourage CNV formation.

## MATERIALS AND METHODS

### Parasite Lines, Compounds, and Treatments

We acquired *Dd2* (MRA-156) and *FCR3* (MRA-731) parasite lines from Bei Resources (ATCC, Manassas, VA). In this study, we were interested in detecting sub-clonal levels of genomic diversity that occur naturally in cell culture (i.e. untreated conditions); therefore, we did not re-clone parasite lines prior to treatment. For low-input genomics, we grew parasites in complete RPMI 1640 with HEPES (Thermo Fisher Scientific, Waltham, MA) supplemented 0.5% Albumax II Lipid-Rich BSA (MilliporeSigma, Burlington, MA) and 50 mg/L hypoxanthine (Thermo Fisher Scientific) and donor A+ human erythrocytes (BioIVT, Hicksville, NY). We grew all cultures at 3% hematocrit at 37°C and individually flushed flasks with 5% oxygen, 5% carbon dioxide, and 90% nitrogen gas. We diluted cultures with uninfected erythrocytes and changed the culture medium every other day to keep parasitemia below 2% during maintenance. We confirmed that all cultures were negative for mycoplasma contamination approximately monthly using a LookOut Mycoplasma PCR detection kit (MilliporeSigma). We synthesized DSM1 as in previous studies (48,49) and purchased aphidicolin (MilliporeSigma).

### Pilot assessments of DSM1 treatment

We conducted multiple independent pilot assessments comparing DSM1 treatment with a replication inhibitor, aphidicolin (see general scheme in **Fig. S1A**). We first acquired high ring-stage cultures (>60%) by synchronizing mixed cultures of parasites twice with 5% sorbitol, ∼48hrs apart. For these pilot assessments, we used sorbitol synchronization to increase the level of ring stage in the cultures. We chose this approach instead of magnetic purification or Percoll gradients in order to limit the time parasites were outside of culture conditions. We then applied 1μM DSM1, solvent control (dimethylsulfoxide, DMSO), or 4.4μM aphidicolin to ring-dominant parasite populations for 12hrs. Following treatment, we washed parasites with sterile 1x phosphate-buffered saline (PBS, Thermo Fisher Scientific), returned them to complete RPMI, and allowed parasites to recover for >36hrs, depending on the experiment (detailed in **Fig. S1B-E**). We tracked parasitemia and parasite viability on an Accuri C6 flow cytometer (BD Biosciences, Franklin Lakes, NJ) as previously performed (42,48,50). At various time points (start at 0h, post-treatment at 12h, or during recovery, **Fig. S1**), we stained parasites with 1x SYBR Green (Thermo Fisher Scientific, stains the parasite nucleus) to assess the proportion of infected erythrocytes (parasitemia) and stage of the parasite development cycle; we also stained parasites with 10nM MitoProbe DiIC1 (5) (Thermo Fisher Scientific, stains active parasite mitochondria) to indicate the proportion of the parasites that were viable after recovery. All pilot experiments were assessed from multiple flasks (>2, **Fig. S1**).

### DSM1 treatment for low input genomics

For parasite treatment for low-input genomics, we synchronized parasites (as above), applied 1μM DSM1 or the DMSO control for 12hrs in duplicate flasks, and allowed recovery for 28-31hrs (**Fig. S2A**). We staggered sample sorting in order to limit the impact on parasite health by removal from culture conditions and allowed parasites to reinvade to produce rings. We removed treatments and tracked parasite number and health as described above (**Fig. S2B-G**). Following reinvasion, we harvested viable 1n ring stage parasites for low-input genomics using flow sorting (details below).

### Parasite Flow Sorting for Low-input Genomics

*Cell sorter calibration & accuracy assessments.* We calibrated the flow sorter (SH800, Sony Biotechnology, San Jose, CA) using the manufacturer’s calibration beads. We accounted for overlaps in the excitation/emission wavelengths using the integrated compensation panel matrix calculation in the SH800 software according to the manufacturer’s procedure. We also manually calibrated the droplet sorting to the nearest 0.2mm, as recommended by the manufacturer, using the 96-well plate setting (Armadillo high-performance 96-well plate, Thermo Fisher Scientific). We evaluated SH800 sorting accuracy prior to low-input harvest using a colorimetric assay as previously described (51). Briefly, we mixed SYBR Green+/MitoProbe+ parasites (see staining details in *Parasite lines, Compounds, & Treatments*) with horseradish peroxidase enzyme (Thermo Fisher Scientific) at a final concentration of 2.5mg/ml. We then sorted parasites into a 96-well plate filled with TMB-ELIZA substrate (Thermo Fisher Scientific) using the single cell (3 drops) instrument setting, in triplicate plates (**Fig. S3A & B**). Formation of a color in the well (blue, green, or yellow) indicates the successful sorting of the enzyme, and therefore parasites, into the well with the substrate. This assessment allowed us to evaluate the accuracy of SH800 sorting (through the evaluation of success for 1-versus 2-cell wells, **Fig. S3C**), the consistency of sorting (through the evaluation of replicates), and the best plate positions for sorting (through the evaluation of performance in different plate rows/columns). Based on these evaluations, we proceeded with isolating 2- and 10-cells per well and avoided sorting into the top 2 rows and the first and last column of the 96-well plate (**Fig. S3D**).

*Parasite isolation & storage*. We stained parasites with SYBR Green and MitoProbe DiIC1 (5) in complete RPMI as above (see staining details in *Parasite Lines, Compounds, & Treatments*), gassed the tubes with 5% CO_2_, 5% 0_2_, 90% N, and placed sample on ice to ensure viability prior to flow sorting of viable, ring-stage parasites (SH800, Sony Biotechnology Inc., San Jose, CA). We used a final concentration of 1 x 10^7^ parasites/ml diluted in sterile 1x PBS (Thermo Fisher Scientific) as input for sorting at the “single-cell setting” (3 drop) into a 96-well plate (Armadillo high performance 96-well plate, Thermo Fisher Scientific) with each well containing 2.375μl of cell lysis buffer (0.025M Tris Ph8.8 (Roche Diagnostics, Indianapolis, IN), 0.01M NaCl (MilliporeSigma), 0.01M KCl (MilliporeSigma), 0.01M (NH_4_)_2_SO_4_ (Thermo Fisher Scientific), 0.001M EDTA (Promega, Madison, WI), and 10% Triton X-100 (MilliporeSigma)). We gated viable 1n ring-stage parasites (**Fig. S4**) and sorted into wells containing cell lysis buffer with an approximate sorting time of 15 min. After sorting, we centrifuged for 30 seconds in a plate centrifuge (MPS1000, Labnet International, Madison, NJ). We immediately overlaid samples with one drop (approx. 25μl) of light mineral oil (BioReagent grade for molecular biology, Millipore Sigma) and sealed the plates with Microamp® Clear Adhesive Film (Applied Biosystems, Waltham, MA) before storage at −80°C until whole genome amplification. All samples were isolated on the same day and stored frozen for ∼1 month before amplification (*FCR3*: 33 days, treated *Dd2*: 35 days, and untreated *Dd2*: 40 days).

### MALBAC Whole Genome Amplification for Low-input Genomics

We handled each plate separately in order to limit cross-contamination during amplification steps. First, we thawed each plate containing sorted parasites (see *Parasite Isolation & Storage*) and added 1mg/ml Proteinase K in sterile 1x PBS (Thermo Fisher Scientific) to a final volume of 2.5μl per well. We heated the plate in a PCR cycler (C1000, Bio-Rad Laboratories, Hercules, CA) at 50°C for 3hrs, followed by 75°C for 20 min and 80°C for 5 min for proteinase k digestion. We amplified parasite genomes using the Multiple Annealing and Looping Based Amplification Cycles (MALBAC) method essentially as previously described ((42), Version 1) with some modifications (Version 2, **Fig. S5**). In summary, 1) we modified the pre-amplification random primer by adding 5 additional degenerate bases with 20% GC-content to increase annealing to AT-rich genome sequences (5’GTGAGTGATGGTTGAGGTAGTGTGGAG<u>NNNNNNNNNN</u>TTT 3’); 2) we performed 19 of the 21 total linear cycles with the *Bsu* DNA Polymerase (Large Fragment, New England Biosciences), which has a lower optimal reaction temperature (37°C) to improve the amplification on AT-rich sequences (52); 3) we lowered the extension temperature from 40/50°C to 37°C during the linear amplification cycles that used the *Bsu* enzyme (see full cycling parameters in **Fig. S5** and impacts of these changes in **Table S1**); and 4) we integrated robotic pipetting (Mosquito LV, SPT Labtech, Melbourn, UK) to increase the throughput of our assays (from 23 samples in Version 1 to 90 samples in the current Version 2) and limit contamination potential.

Overall, we performed 21 total linear cycles (19 cycles with *Bsu* polymerase and 2 cycles with *Bst* polymerase, New England Biolabs, Ipswitch, MA) and 17 total exponential amplification cycles using Herculase II Fusion DNA polymerase (Agilent Technologies, Santa Clara, CA). During amplification steps, we employed standard steps to limit contamination (42). For automated pipetting of the enzyme solution during linear cycles, tips were changed after each round of pipetting. Post-amplification, we purified amplified DNA with Zymo DNA Clean & Concentrator-5columns (Zymo Research, Irvine, CA) according to the protocol and ran 2µl of all samples on 1% agarose gels to check for the presence of DNA (generally, if >30ng/µl, samples could be visualized with a size range of 100 to >1500bp).

### Assessments of Amplification Success for Low-input Genomics

*DNA quantification.* We quantified the MALBAC-amplified DNA using a Qubit fluorimeter (Qubit 1X dsDNA High Sensitivity Assay Kit, Thermo Fisher Scientific) (**Fig. S6**).

*Droplet Digital PCR.* To confirm the presence of parasite DNA in MALBAC-amplified samples, we performed droplet digital PCR (ddPCR) as described previously using the QX2000 droplet generator, C1000 thermocycler, and QX2000 droplet reader (Bio-Rad Laboratories) (48,53) (**Fig. S7**). We used duplex assays to evaluate two parasite genomic loci concurrently (*pfmdr1:* Forward-TGCCCACAGAATTGCATCTA; Reverse-ACCCTGATCGAAATGGAACCT; Probe - TCGTGTGTTCCATGTGACTG; *pfhsp70:* Forward-TGCTGTCATTACCGTTCCAG; Reverse - AGATGCTGGTACAATTGCAGGA; Probe -AGCAGCTGCAGTAGGTTCATT (Integrated DNA technologies, Newark, NJ). The reaction master mix contained 600nm of forward and reverse primers, 50nm probes, 10µl of ddPCR Supermix for Probes (2x, Bio-Rad Laboratories), 3µl of nuclease-free water (QIAGEN), and 1.5ng (5µl) of template DNA per assay (total of 20µl). We used the following cycling conditions for PCR amplification: 10 min at 95°C initial denaturation step, 1 min at 95°C second denaturation step, and 2 min at 58°C annealing and extension step (ramp rate of 1°C per second), the second denaturation step and the annealing/extension step repeated 60 times, and then 10 min at 98°C to halt the reaction (53). In addition to running amplified samples to assess amplification success, we ran ddPCR with bulk genomic DNA as a positive control, no template controls (water replaced DNA), and material from “no cell” wells to assess cross-well contamination. We considered the samples positive for parasite DNA if there were more than 50 total positive droplets in target-positive clusters. For comparisons to DNA amounts (**Fig. S6**), we plotted total nanograms against total positive droplets (from both *pfmdr1* and *pfhsp70* loci) and performed a simple linear regression in PRISM (GraphPad, La Jolla, CA).

*High Resolution Melting Assay.* To assess potential contamination between MALBAC-amplified samples, we performed asymmetric PCR amplification of the *pfdhps* locus followed by high-resolution melting (HRM) as described previously (54,55) (**Fig. S8**). The *pfdhps* locus at codon 613 is distinct in *Dd2* and *FCR3* parasite lines (*Dd2*: Ser-613 and *FCR3*: Ala-613, (56)). Each 20µL reaction contained 8µl of the 2.5x LightScanner Master mix (BioFire^TM^ Defense, Salt Lake City, Utah, USA), 1/10µM of forward/reverse primers and 8µM probes targeting the *pfdhps* gene position 613: Forward - CTCTTACAAAATATACATGTATATGATGAGTATCCACTT; Reverse-CATGTAATTTTTGTTGTGTATTTATTACAACATTTTGA; Probe - AAGATTTATTGCCCATTGCATGA/3SpC3, (Integrated DNA technologies), 7µl of nuclease free water, and 3µl of DNA (∼0.05ng total). We used the following cycling conditions for PCR amplification with the Rotor-Gene Q instrument with a 72-well rotor (QIAGEN): 95°C for 5 min, 45 cycles of 95°C for 10s, 55°C for 30s, and 72°C for 10s, followed by a pre-melt at 55°C for 90s, and a HRM ramp from 65°C to 90°C, with an increase of 0.1°C every 2s. We plotted the change in fluorescence versus temperature (dF/T) using Rotor-Gene Q software (version 2.3.5, build 1; QIAGEN) and compared HRM peaks of amplified samples to bulk genomic DNA and plasmid controls.

### Bulk DNA Extraction for Short-Read Sequencing

We extracted bulk DNA for short-read sequencing as previously performed (42). Briefly, we lysed erythrocytes with 1.5% saponin and washed the parasite pellet 3 times with 1x PBS (Thermo Fisher Scientific), before resuspension in a buffered solution (150mM NaCl (MilliporeSigma), 10mM EDTA (Promega Corporation, Madison, WI), and 50mM Tris pH7.5 (Roche Diagnostics)) to a total volume of 500µl. We then lysed the parasites with 10% sarkosyl (MilliporeSigma) and 20mg/ml proteainase K (Thermo Fisher Scientific) at 37°C overnight before DNA purification using standard phenol/chloroform/isoamyl alcohol extraction and chloroform washing steps (2 times each, (42)). Finally, we precipitated DNA using 100% ethanol with 100mM of sodium acetate overnight in DNA-lo bind tubes (Eppendorf, Enfield, CT) and then washed twice with 70% ethanol before resuspension in 50µl nuclease free water (QIAGEN). We stored bulk genomic DNA at −20°C until sequencing library preparation.

### Low-input Genomics Sample Selection & Short-Read Sequencing

*Low-input sample selection.* We selected 16 low-input samples from the *FCR3* plate, 36 samples from the untreated *Dd2* plate, and 38 samples from the treated *Dd2* plate for short-read Illumina sequencing. We based our selection on the quantity of the MALBAC amplified DNA and presence of parasite DNA using ddPCR (**Figs. S6 & S7**). In summary, the majority of *FCR3* and untreated *Dd2* samples yielded quantifiable parasite DNA following MALBAC amplification (53/60 *FCR3* samples and 60/60 untreated *Dd2* samples); in these conditions, we chose samples randomly to proceed with sequencing (indicated in **Fig. S6**). For treated *Dd2* samples, we chose samples for sequencing if they had adequate DNA quantity (>10ng total, 30 samples, **Fig. S6**) or had ddPCR results showing the presence of parasite DNA (an additional 8 samples).

*Short-read sequencing.* Before short-read sequencing, we sheared bulk samples and low-input samples using Covaris M220 Focused Ultrasonicator for 150s and 130s, respectively, to generate fragment sizes of ∼350bp as evaluated by an Agilent 2100 Bioanalyzer using the High Sensitivity DNA kit (Agilent, Santa Clara). We adjusted the volume of sheared samples with nuclease-free water up to 50µl. For samples with >100ng (**Table S2**, including bulk and MALBAC amplified samples), we diluted them to 1.2-2ng/µl; for samples <100ng, we proceeded with no dilution. We used NEBNext Ultra II kit (Illumina Inc., San Diego, CA) to prepare libraries for sequencing with 3 cycles of PCR amplification, as performed previously (42). We quantified the resulting libraries using NEBNext Library Quant Kit (Illumina Inc.) before sequencing on the Illumina Nextseq 550 using 150 bp paired-end cycles.

### Short-Read Sequence Processing & Analysis

*Read processing and alignment*. We performed short-read quality control steps as described previously (40,42). Briefly, we reordered and removed singletons and subsequently interleaved paired reads using BBMap, filtered for high-quality reads (>Q30), trimmed the MALBAC common sequence, PhiX, and Illumina adapters from the remaining reads with the BBDuk tool within BBMap, and aligned reads to the *pf3D7-62_v3* reference genome using the Speedseq genome aligner (57). We removed reads that mapped to VAR regions from bam files according to previously defined genomic coordinates (58). We filtered out reads with low mapping quality (mapq<30) and duplicated reads using SAMtools (59). We calculated the percentage of reads that map to the *P. falciparum* genome by dividing the number of mapq>30 mapped reads by the number of total unaligned reads (mapped reads/total reads). We employed Qualimap to report mean coverage and standard deviation across the genome (60). Using non-overlapping 20kb size bins, we calculated the coefficient of variation of read coverage by dividing the standard deviation of coverage within a bin by the mean across a sample and multiplying by 100 (61,62) (R version 4.4.2). Short-read data is available at NCBI Sequence Reads Archives under project PRJNA1201106.

*BLAST-based evaluation of read composition.* We blasted reads from low-input genomics samples essentially as previously described (42). We also performed an additional BLAST analysis to understand read composition. We randomly downsampled and extracted 5,000 reads from either 1) all aligned reads (mapq>1) or 2) unmapped reads (mapq>30, **Fig. S9**). To ensure that BLAST alignments consisted of high-quality reads, we manually filtered out low complexity (%N>50) and short sequences (length < 100bp). We used these reads as input for a custom pipeline, where we blasted reads against a database consisting of common genomes downloaded from NCBI (i.e., human, mouse, mycoplasma, bacterial genera, etc). For the BLAST step, we employed a Conda installable BLASTN v2.16.0+ for analyzing the reads (63). In order to count as a hit, the sequence must have 90% of its bases aligned and have an e-value of <e-20. The pipeline is designed to return the top 10 blast hits with these criteria, and either call the most significant hit (lowest e-value), or if alignments from the *Plasmodium* genus were present in the result (but not the most significant hit), call the most significant *Plasmodium* hit. We included this step to account for possible mismatching due to the abundance of highly repetitive sequences in the *Plasmodium* genome. We also used a custom script to quantify the abundance of hits based on the genus related to each hit’s accession number. If the pipeline identified a hit, but the genus was not represented in the custom database, the hit was called ‘unknown’. The code for the pipeline, including construction of the database, is available at https://doi.org/10.6084/m9.figshare.29885879.

*Single nucleotide polymorphism analysis*. We performed SNP genotyping and analysis as previously (42), based on the MalariaGen *P. falciparum* Community Project V6.0 pipeline (64–67) using the *pf3D7-62_v3* reference genome (**Fig. S10**). Briefly, we applied GATK’s Base Quality Score Recalibration using default settings. We detected potential SNPs using GATK’s HaplotypeCaller and subsequently genotyped the SNPs using CombineGVCFs and GenotypeGVCFs. We employed GATK’s VariantRecalibrator using previously validated SNP datasets (68) and applied GATK’s ApplyRecalibration to assign VQSLOD scores (67). We filtered the resulting SNPs for those with VQSLOD > 6 and for a GT quality metric >20 to ensure high-quality variant calling. We only selected variants flagged as Bi-allelic to simplify the analysis. For SNP Principal Component Analysis (PCA), we merged experiment-wide SNP data (described above) into a single file. Then we merged the VCF into a large matrix and converted the genomic data into numeric information using the ‘vcfr’ package in R (Version 4.2.3) (https://CRAN.R-project.org/package=vcfR). We excluded individual SNPs if >25% of the samples lacked a call in this position or if all calls were the same for each sample in that position). We scaled the remaining SNPs around the origin using the ‘scale’ R function. We then calculated the principal components using the ‘prcomp’ R function and scored the dataset using the ‘scores’ function from the ‘vegan’ R package (https://CRAN.R-project.org/package=vegan).

### Copy Number Variation Analysis

*CNV calling in bulk samples.* We performed CNV detection for bulk samples similar to as previously described (40,42). Briefly, we called CNVs independently using two methods, CNVnator (read depth based calling, (69)) and LUMPY (split and discordant read based calling, (70)). To Identify CNVs called in both methods, we used SVCROWS to define overlapping CNV regions relative to their size (43). Briefly, SVCROWS uses a reciprocal-overlap-based approach (i.e. two CNVs must be overlapping each other at, or greater than, a defined threshold) to determine if two CNVs are close enough in their genomic position to be called the same. The program utilizes adaptive thresholds for overlap based on the sizes of the CNVs being compared, ensuring that we account for shifts in CNV position, particularly in regions of high variance. For known CNV calls, we used the following SVCROWS input parameters: ExpandRORegion = FALSE, BPfactor = TRUE, DefaultSizes = FALSE, xs = 3000, xl = 10,000, y1s = 300, y1l = 1000, y2s = 50% and y2l = 80%; based on the average size of a *P. falciparum* gene (∼2.3kb) and intergenic region (∼2kb). Similar to our previous study (42), we identified 3 known CNVs that were called by both LUMPY and CNVnator methods in the core genome of bulk samples (untreated and treated). We determined known CNV boundaries using SVCROWS: *pfmdr1* (*Pf3D7_05_v3*, 888001-970000, 82kb), *pf11-1* (*Pf3D7_10_v3*, 1521345-1541576, 20kb), and *pf332* (*Pf3D7_11_v3*, 1950201-1962400, 12kb).

*Downsampling.* For CNV analysis that assessed downsampled data, we first converted the processed bam files back into FASTQ files using SAMtools and then used the reformat.sh option of BBtools to select 1.3M reads from each FASTQ (represented the fewest number of reads from a sample that passed quality filtering from the final dataset). We then realigned files to the reference genome (*pf3D7-62_v3*).

*CNV calling in 2-cell samples.* We employed two methods for CNV detection in the core genome of low-input samples; LUMPY is a split/discordant read strategy with high sensitivity (70), and HapCNV is a read coverage-based strategy designed for haploid genomes (44). We ran LUMPY as part of Speedseq with default parameters as previously described (40). We filtered resulting structural variants to include only duplications (DUP, >1 copy of a region) and deletions (DEL, one less copy of region than the reference). We then filtered those calls for those GQ > 20 to ensure high-fidelity calls. In HapCNV, we used a quality control and bias correction procedure to exclude bins of poor quality and remove bias introduced by GC content and mappability variation. We then constructed a pseudo-reference for each *Dd2* low-input sample using within-*Dd2* information, which enabled control of background noise while preserving CNV signals after normalization. Finally, we used a circular binary segmentation algorithm (CBS, (71)) to detect copy number change points followed by a Gaussian Mixture Model (GMM, (72)) for CNV identification.

For statistics, we used PRISM (GraphPad, La Jolla, CA), using unpaired parametric T-tests with Welch’s correction. We calculated standard error in Microsoft Excel.

*Defining CNV regions/determination of “rarity” in CNV calling.* Small differences in sequence quality surrounding a read can lead to shifted breakpoint determination for biologically identical CNVs, which is especially true for low-input genomics datasets (28). To account for this, we used SVCROWS to determine whether two CNV signals were the same within and between samples (see *CNV calling in bulk samples* for input parameters (43)). We assigned the categorizations of “rare” and “common” by assessing the CNV region frequency within datasets as performed in other studies (44,73). “Rare” CNVs were defined as occurring in <10% of the samples within a treatment group; “common” CNVs were defined as occurring in ≥10% of samples within a treatment group; “known” CNVs were defined by CNVs called in bulk samples (see above, *CNV calling in bulk samples*).

*High-confidence CNV region identification.* To identify “high-confidence” CNV regions called by both HapCNV and LUMPY methods, we compared the ‘consensus list’ generated by SVCROWS (43) for each detection method by combining them into a single input file. Because there is a large disparity of average CNV region lengths between the two methods (HapCNV = ∼40kb, LUMPY = ∼4.3kb), we relaxed the stringency of the SVCROWS parameters for this analysis. Our input parameters to generate the “high-confidence” list were as follows: xs = 3000, xl = 6000, y1s = 500, y1l = 1500, y2s = 30% and y2l = 60%. We defined high-confidence CNV regions as those that had >1 match from both HapCNV and LUMPY that was the same type (i.e. either duplication or deletion or mixed). For Venn diagram generation, we calculated overlaps using SVCROWS “Scavenge” mode (43), input parameters: ExpandRORegion = FALSE, BPfactor = TRUE, DefaultSizes = FALSE, xs = 3000, xl = 6000, y1s = 500, y1l = 1500, y2s = 30% and y2l = 60%). We systematically compared lists for each overlap comparison, and if regions had at least one match in an opposing dataset, we considered it a match. We used the *draw.quad.venn* function in the “VennDiagram” R package (R 4.2.1) to generate the diagram.

*Breakpoint analysis of high-confidence CNV regions.* We assessed features of breakpoints flanking high-confidence CNVs as performed previously (40). Briefly, we used the CNV breakpoint coordinates either set by LUMPY or HapCNV to extract a 2000bp window around the 5’ or 3’ CNV end (±1000bp from the breakpoint). Although Huckaby et al. solely used LUMPY to determine breakpoints (40), we employed CNV boundary information from both CNV callers (LUMPY and HAPCNV) merged using the Expand-RO function in SVCROWS to construct the conserved CNV region ((43), see parameters in *High-confidence CNV region identification*). In this analysis, breakpoints are called using either and/or both HapCNV and LUMPY boundaries on either end, which captures variance but can limit accuracy. For this reason, we only identified breakpoints of high-confidence CNVs, which are conserved between CNV-callers. With these defined breakpoints, we then used Vienna 2.1.9 folding prediction software with Mathews 2004 DNA folding parameters, wherein G-quadruplexes, GU pairing, and lonely base pairs were disallowed to predict stable structures in that region based on Gibbs free energy estimates in sliding 50bp windows. We called stable hairpins by determining local minima of Gibbs free energy across the 2000bp region using the same 50bp windows. We reported the position of the stable structure closest to the breakpoint using the threshold of −5.8kCal/mol, which we previously reported as the cutoff for stable secondary structure formation in this genome. If no stable hairpins could be determined within 2kb around the breakpoint, we recorded that breakpoint as “unresolved”. We also calculated the GC content of the breakpoints from the 100bp window surrounding the stable hairpin, whether this position was genic or intergenic, and the size of the resulting CNV as the distance between the two breakpoints.

*Gene Ontology enrichment and protein class identification from high-confidence CNV regions.* We used the online Gene Ontology Resource (geneontology.org) to perform GO enrichment analysis using the PANTHER Classification System (74,75). Since a large portion of the *P. falciparum* genome remains unannotated (PlasmoDB, ∼30%) and the majority of molecular functions remain unclassified (92.8%), we used the Panther Protein class assessment (version 19.0, only 55.9% remained unclassified) with default statistics (Fisher’s test with FDR adjusted p value of <0.05 for significance, which is recommended for small counts and overlaps between classes). We used the web tool to represent protein classes on pie charts.

*Clinical importance of high-confidence CNV regions.* We assessed the clinical importance of genes within high-confidence CNVs by first collecting gene IDs from high-confidence duplications and deletions. To generate this gene list, we used SVCROWS “Hunt” mode ((43), input parameters: BPfactor = TRUE, DefaultSizes = FALSE, xs = 3000, xl = 6000, y1s = 300, y1l = 600, y2s = 30, y2l = 60), which takes a secondary input list of known genes (Pf3D7_62_v3, Plasmodb.org) and overlaps CNVs to the gene list using a size-weighted scale. We then performed a literature search to identify genes within the list that are essential for erythrocytic stage, mosquito stage (including gametocyte, gamete, ookinete, sporozoite formation), or liver stage *P. falciparum* or *P. berghei* development. We also included genes that produced known parasite antigens important for immune recognition/vaccine escape or contributed to clinical drug resistance. We noted the Mutagenesis Index Score (MIS) for each gene, which estimates essentiality in erythrocytic stage parasites (76), from the PlasmoDB database (77). To assess overlap of high-confidence CNVs with a large catalogue of *P. falciparum* CNVs from clinical isolates, we collected the gene list from high-frequency CNVs (>1%, >300bp) identified in clinical isolates (Supplementary Table 3 from (38)) and manually reformatted the list to match SVCROWS input style guidelines (43). We then assessed how often these genes overlapped with our high-confidence gene list using the SVCROWS “Hunt” mode described above.

## RESULTS

### Refined low-input genomics pipeline increased efficiency and accuracy

We developed a robust pipeline for assessing the frequency of de novo CNVs in the *P. falciparum* genome. We adapted our single cell genomics method to improve parasite isolation and whole genome amplification steps over our previous study (42). In this modified protocol, we used fluorescence-activated cell sorting (FACS) to isolate viable, 1n parasites to improve efficiency and modified aspects of our whole genome amplification method to improve coverage (**Table S1**, **Fig. S5**). We also sorted low-cell populations to increase accuracy (e.g. 2-cells per well, **Fig. S3**), and added quality control PCR-based steps that confirmed our samples were of high quality prior to short-read sequencing (**Fig. S7 & S8**). The resulting low-input genomics pipeline consisted of basic steps including parasite isolation using flow sorting, whole genome amplification using a modified MALBAC-based approach, quality control confirmation, short-read sequencing, and CNV analysis (**Fig. 1B**).

### Replication stress followed by a recovery period led to isolation of viable parasites

To explore the impact of antimalarial stress on de novo CNV generation, we treated the parasites with a compound that targets *Plasmodium* dihydroorotate dehydrogenase (DSM1, (80)). The application of DSM1 for an extended period kills parasites (>48hrs at 10x the EC_50_, (81)). However, we found that when applied to ring-stage parasites for a short period of time (see general timeline in **Fig. 1C**), DSM1 stalls parasite progression through the life cycle in a reproducible and reversible manner (**Fig. S1B, C & E**); this effect was similar to parasite treatment with the replication inhibitor, aphidicolin ((82,83), **Fig. S1D & E**). While aphidicolin stalls replication through inhibition of B-family DNA polymerases (83,84), DSM1 likely impacts replication through depletion of pyrimidine pools as is observed for other forms of dNTP depletion (85,86). For this reason, we now refer to DSM1 treatment as “replication stress”.

For the low-input genomics harvest, we applied DSM1 to ring-stage parasites for 12h (**Fig. 1C & S2A**). Similar to our pilot experiments (**Fig. S1B, C & E**), we observed that DSM1-treated parasites exhibited a slightly decreased growth rate and altered stage progression compared to untreated parasites (**Figs. 1D & S2B-G**). We harvested viable parasites after a recovery period (an additional ∼30hrs, **Fig. 1E**, or ∼42 hours from the treatment start, **Fig. 1C**), where we allowed parasites to complete an additional round of replication and erythrocyte invasion to produce those that have a single, haploid genome (ring stage, **Fig. S2E, G**).

As a control for cross-sample contamination between isolation wells, we isolated untreated parasites with different genetic backgrounds (*FCR3* vs *Dd2*, **Table 1**, **Fig. 1D & E**). Prior to isolation of low-cell populations, we confirmed that *FCR3*, untreated *Dd2*, and DSM1-treated *Dd2* parasites were at a similar life cycle stage and viability (**Table 1**); we saved a portion of these samples for parasite population sequencing (i.e. bulk samples). We then proceeded to isolate small populations of viable, ring stage parasites using FACS (**Fig. S4**) for whole genome amplification and sequencing.

**Table 1:** Parasite density, staging, and health at isolation for low-input genomics.

| Line/Treatment* | Mean % Parasitemia<sup>§</sup> | Mean % Rings** | Mean % Viability<sup>@</sup> |
| --- | --- | --- | --- |
| Untreated *FCR3*<sup>^</sup> | 0.6% | 90% | 91% |
| Untreated *Dd2*<sup>^</sup> | 0.8% | 87%<sup>&</sup> | 94% |
| DSM1-treated *Dd2* | 0.6% | 77%<sup>&</sup> | 95% |

\*Treatment conditions: 1 μM DSM1 (~10x EC<sub>50</sub>) for 12hrs prior to 31hrs of recovery. Untreated samples were incubated with DMSO as a solvent control for 12hrs and allowed to recover for 28-30hrs (**Fig. 1E**). <sup>§</sup>Parasitemia was determined by calculating the number of infected erythrocytes (SYBR Green+) compared to uninfected erythrocytes (SYBR Green-) (**Fig. S2C-G**). \*\*Mean % Rings value was collected from “post-recovery” plots in **Fig. S2E/G** (R1 SYBR green plots). <sup>@</sup>Viability of parasites was determined by measuring the mitochondrial membrane potential (Mitoprobe+) and calculating % of total SYBR Green-based parasitemia (plots presented in **Fig. S2E, S2G, and Fig. S4**). <sup>^</sup>*FCR3* (Africa) and *Dd2* (Southeast Asia) parasite lines are from distinct geographic origins (87,88). <sup>&</sup>The initial % rings were 91.8% for both untreated and treated conditions (**Fig. S2C**).

### Quality assessments showed effective isolation and amplification of low-input samples

We sorted ring-stage low-cell populations from each parasite group into 60 wells of a 96-well plate (*FCR3*, untreated *Dd2,* and treated *Dd2*), including 6 wells with 10-cells and 56 wells with 2-cells. Ten-cell wells served as positive controls for the whole genome amplification step and 2-cell wells provided the optimal balance between sorting accuracy (**Fig. S3**) and de novo CNV detection. Zero cells were sorted into the top 2 rows of the plate (“no-cell” wells). After parasite lysis and applying *Pf*MALBAC version-2 whole genome amplification (**Fig. S5**), we assessed amplification success using three approaches. First, we measured the resulting DNA quantity across 80% of the amplified wells (**Fig. S6**). On average, MALBAC amplification in wells that contained sorted parasites yielded ∼120ng of total DNA per reaction, with a ∼10% increase in DNA for 10-vs 2-cell samples (mean of 127ng versus 116ng total, respectively). We did not detect position-based bias across plates or appreciable amplification from no-cell wells, but we observed that treated *Dd2* wells had ∼3-fold lower levels of amplification than other samples (mean of 51ng versus 151ng total, respectively). Of note, there was little difference in mean amplified DNA amounts between the two untreated sample groups (*FCR3* at 146ng and untreated *Dd2* at 152ng).

Second, we performed droplet digital PCR (ddPCR) for parasite-specific genes on approximately one-third of wells to confirm the amplification of the parasite genome. DdPCR for *pfmdr1* and *pfhsp70* displayed that wells with measurable DNA contained amplified parasite DNA (**Fig. S6** and **S7**). Additionally, we confirmed that 2-cell wells with very low total DNA amounts (**Fig. S6**) were positive for parasite genomes while “no-cell” wells did not show evidence of parasite material (**Fig. S7A-C**). We observed a significant correlation between total DNA and positive ddPCR droplet counts (black dotted line, p value of 0.0004) but also identified some samples where DNA quantity was low (<50ng) when measured using Qubit but parasite-specific signal was high (red points, all treated *Dd2*, **Fig. S7D**). This observation indicated that DNA quantification is not the best way to measure amplification success; target-based methods like ddPCR are more helpful to estimate the quantity of amplified material.

Finally, we employed high-resolution melting analysis to profile a drug resistance marker that differs between *FCR3* and *Dd2* parasites (56). By assessing the *pfdhps* SNP profile of amplified genomes and comparing it to the parental profile in ∼10% of samples, we confirmed that there was no evidence of cross-sample contamination during the preparation and amplification steps (**Fig. S8**). Therefore, we proceeded to sequence the amplified bulk and low-input samples (**Table S2**).

### Coverage deviation and SNP profiles exhibited expected trends in low-input samples

We sequenced 3 bulk samples and 90 low-input samples using Illumina short-read sequencing (**Table 2** and **S3**). Overall, sequencing proceeded well as indicated by coverage and coverage deviation of the bulk samples, as well as an equivalent mean mapping quality across all samples (**Table 2**). Because we noticed that treated *Dd2* wells had lower levels of DNA following amplification (**Fig. S6**), we sequenced higher amounts of material for this condition; this choice impacted mean coverage levels where treated *Dd2* samples had ∼4-times higher coverage than untreated *Dd2* samples (**Table 2**). As expected, based on previous studies (42), coverage deviation was ∼3-fold higher in low-input samples when compared to bulk samples, reflecting the bias of the whole genome amplification step to over-or under-amplify specific genomic regions.

**Table 2:** Sequencing Summary for low-input samples and paired bulk samples.

|  |  | No. of samples<br>** | Mean no. mapped reads per sample | Mean coverage per sample | Mean Coefficient of Variation (CV#) | Mean mapping quality |
| --- | --- | --- | --- | --- | --- | --- |
| Bulk | Untreated *FCR3* | 1 | 9,966,137 | 54.9 | 59.5 | 58.2 |
| Low-input | Untreated *FCR3* | 16 | 2,694,209 | 13.7 | 105.1 | 58.2 |
| Bulk | Untreated *Dd2* | 1 | 5,735,107 | 33.6 | 33.3 | 58.5 |
| Low-input | Untreated *Dd2* | 33 | 2,579,674 | 13.2 | 82.6 | 58.3 |
| Bulk | Treated *Dd2* | 1 | 72,380,215 | 416.3 | 35.2 | 58.5 |
| Low-input | Treated *Dd2* * | 36 | 9,936,903 | 53.7 | 86.5 | 58.4 |

\*Four times more material was loaded on the flow cell for the treated samples than the untreated samples due to lower initial amplification yields (Fig. S6).
\*\*Excludes samples that were removed due to low coverage. For low-input samples, analysis includes both 10- and 2-cell samples.
#CV is the coefficient of variation of normalized read abundance as in (42)

The percentage of high-quality reads that mapped to the *P. falciparum* genome was high across all samples (mean of 67% of all reads). To identify any contaminating reads, we also blasted a subset of high-quality reads (%N <50, length >100bp, and read q score >30) from each sample to a database of genomes from other organisms. While the majority yielded matches to the *P. falciparum* genome (97.5%), a very small number of high-quality reads matched sequences from different organisms (**Fig. S9**). Together, this information shows efficient amplification of the parasite genome and little contribution of environmental contamination during sample preparation. We removed five low-input samples from further analysis based on low coverage levels; on average, excluded samples had ∼7-times lower coverage than other low-input samples (**Table S3**). Of the remaining samples, 18 were 10-cell samples and 57 were 2-cell samples. Although mean coverage was ∼2-fold higher for 2-cell samples due to the higher coverage of treated low-input samples, the mean normalized deviation was similar between 10- and 2-cell samples (3.5 and 3.1, respectively).

To once again check for cross-sample contamination, we tracked SNPs in the low-input samples compared to bulk samples. Despite some variation due to the non-clonal nature of parasite lines (see *Materials and Methods*), low-input SNP profiles were similar to their corresponding bulk sample as compared using PCA (**Fig. S10A**). After normalizing total SNPs to mapped reads, we detected a lower rate of SNPs in treated samples compared to untreated counterparts (p value of 0.0001, **Fig. S10B**). While we did not detect a correlation between normalized total SNPs and amplification quality (**Fig. S10C**, R_2_: untreated *Dd2*, 0.01; treated *Dd2,* 0.01), we did observe a positive correlation between SNP number and coverage depth in treated versus untreated *Dd2* samples (**Fig. S10D**, R_2_: untreated *Dd2*, 0.66; treated *Dd2,* 0.27). This latter finding indicates that the difference in SNP numbers between sample groups is likely due to varying sensitivity at different levels of read coverage (89,90).

### Experimental and computational advances improved known CNV calls across low-input samples

For the current study, we employed two different CNV calling methods in low-input samples. HapCNV is a novel read coverage-based CNV calling method specifically designed for low-cell data from haploid genomes (44). In contrast to traditional methods that arbitrarily select reference samples for CNV data normalization, HapCNV constructs a genomic location (or bin)-based pseudo-reference as a comparison baseline. This step systematically alleviates amplification bias for the identification of de novo CNVs. On the other hand, LUMPY is an established CNV calling method that exhibits high sensitivity due to the incorporation of multiple CNV signals (i.e. split and discordant reads) generated from short-read sequencing. It is particularly well-suited for detecting low-frequency variants; however, as for many CNV callers, high sensitivity leads to higher false positives (70,91,92).

Using these two distinct CNV calling methods, combined with a recently developed CNV counting approach (SVCROWS (43)), we evaluated the presence of known CNVs in our 2-cell samples (**Fig. 2A & B**). The identification of known CNVs (i.e. those identified in the bulk sample, see *Materials and Methods*) within low-input samples displays the utility of the specific CNV calling methods for different size CNVs in specific genome locations. In our previous study, we identified 2 of the 3 known CNVs in ∼10% of single cell genomes (42). In the current study, we identified the *pfmdr1* amplicon in 100% of 2-cell samples using HapCNV (57/57) and 79% of samples using LUMPY (45/57). We did not identify the *pf11-1* amplicon in any 2-cell samples using HapCNV (0/57) but detected this locus in 75% of samples using LUMPY (43/57). Finally, we identified *pf332* CNVs in 46% of 2-cell samples using HapCNV (26/57) and 100% of samples using LUMPY (57/57). Although our two studies used different CNV calling methods and are not directly comparable, the overall improvement in the detection of known CNVs in the current study is likely due to advances in both the whole genome amplification method to limit amplification bias (**Table S1**) and recently developed analysis approaches. When we evaluated the detection of three known CNVs specifically in *Dd2* low-input samples, we observed a somewhat higher rate of known CNV detection by either method in treated samples (**Fig. 2B**, HapCNV: mean of 1.2 out of 3 total CNVs for untreated and 1.7 for treated samples (increase of 42%), LUMPY: mean of 2.1 out of 3 total CNVs for untreated and 2.9 for treated samples (increase of 38%, **Table S4**).

**Figure 2.**
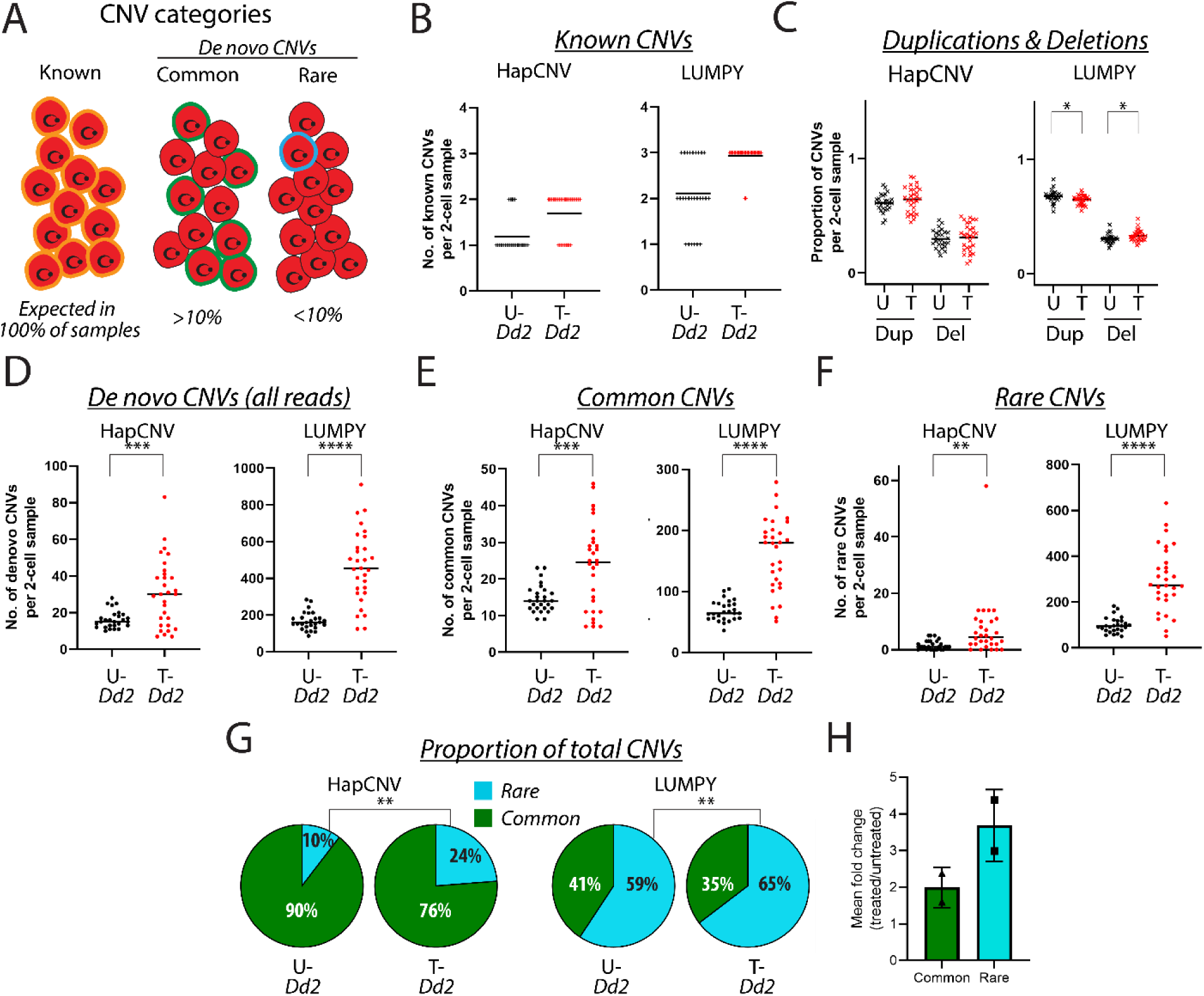
**Low cell genomics displays an increase in de novo CNVs following replication stress**. Number of CNVs from untreated (U-*Dd2*) and treated (T-*Dd2*) 2-cell samples from two CNV analysis methods: HapCNV and LUMPY. Statistics for all plots use an unpaired T-test with two tailed Welch’s correction (****:p value <0.0001; ***: <0.001; **: <0.01; *:<0.05; no stars: not significant). Analyses include all reads. Line at mean value for each dataset. **A**. Depiction of CNV categories used in the analysis. Known CNVs (orange) are detected in bulk samples and present in all low cell samples. De novo CNVs are not present in bulk samples are considered common (green, >10%) or rare (teal, <10%) depending on their frequency across the 2-cell samples. **B**. Detection of three known CNVs in 2-cell samples. Known CNVs were identified in *Dd2* bulk sequence (either *pfmdr1, pf11-1, or pf332*). 0: no known CNVs were detected in 2-cell sample; 1/2/3: one/two/or three known CNVs were detected in 2-cell sample (see **Table S4** for sample counts). **C.** Proportion of total CNVs detected as duplications (Dup) or deletions (Del) in untreated (U) or treated (T) 2-cell *Dd2* samples (p values: 0.02 for Dup and 0.03 for Del from LUMPY). **D**. Detection of de novo CNVs (common and rare combined) from all reads (p values: 0.0002 for HapCNV, <0.0001 for LUMPY). **E**. Detection of common CNVs from all reads (p values: 0.0004 for HapCNV, <0.0001 for LUMPY). **F**. Detection of rare CNVs from all reads (p values: 0.008 for HapCNV, <0.0001 for LUMPY). **G**. Proportion of total CNVs detected as rare and common from all reads; pie charts show the mean but statistics are calculated using all data points from the rare CNV category (p values: 0.003 for HapCNV, 0.009 for LUMPY). Pie chart size does not represent total de novo CNV numbers (∼12x higher for LUMPY, **Table S4**)**. H**. Mean fold change between rare and common CNVs detected by HapCNV and LUMPY in untreated and treated *Dd2* 2-cell samples (see **Table 3**).

**Table 3.** Mean CNV counts per 2-cell sample using two analysis methods.

| CNV detection method<sup>§</sup> | Condition (2-cell only) | Rare CNVs per sample (SE)* | Common CNVs per sample (SE)* | Combined de novo CNVs per sample (SE)^ |
| --- | --- | --- | --- | --- |
| HapCNV | treated *Dd2* | 7 (1.9) | 23 (2.1) | 15 (3.3) |
|  | untreated *Dd2* | 1.7 (0.4) | 15 (0.8) | 8 (1.0) |
|  | fold change | **4.4** | **1.6** | **1.9** |
| LUMPY | treated *Dd2* | 297 (25) | 163 (11) | 230 (35) |
|  | untreated *Dd2* | 99 (6.5) | 68 (3.1) | 84 (9.3) |
|  | fold change | **3.0** | **2.4** | **2.7** |

\*Includes both duplications and deletions. Values are calculated by taking the mean of CNV counts per sample within each category. ^De novo CNV counts combined the subcategories of rare (<10% of samples) and common CNVs (>10% of samples, absent in bulk). §CNV analysis performed using all reads (downsampled data presented in **Fig. S12**).

### De novo CNVs in low-input samples consisted of rare and common CNVs

We next sought to quantify de novo CNVs in low-input samples. We defined de novo CNVs as those that are not present in the bulk sample and categorized them based on their frequency in low cell samples. As defined in other studies (44,73), “common” CNVs were present in a larger number of genomes (≥10% of the same sample type, i.e. untreated or treated), and “rare” CNVs were those that occurred in a small proportion of samples (<10% of the sample type) (**Fig. 2A**). Overall, LUMPY detected more total de novo CNVs than HapCNV across all samples (∼12-fold), and the proportion of rare versus common CNVs varied depending on the method (18% vs 82% for HapCNV, 61% vs 39% for LUMPY, respectively, **Table S4**). Additionally, de novo CNVs were more often identified as duplications than deletions for both CNV calling methods (**Fig. 2C**).

To understand the nature of common and rare CNVs, we also assessed how often their locations overlapped across the two sample types (untreated and treated, **Fig. S11**). This analysis is useful for tracking common/rare category utility and relevance. For utility, this step acts as a sanity check since, by definition, we do not expect rare CNV locations to overlap as often as common CNVs. For relevance, de novo CNVs with conserved locations across sample types are less likely to represent true CNVs newly arising in a genome. As expected, we identified common CNVs with conserved locations between untreated and treated samples (22% for HapCNV and 52% for LUMPY, **Fig. S11A**). The lower rate of overlapping calls across samples for HapCNV is likely due to the bin-based normalization strategy to remove amplification artifacts (44). Conversely, we found that rare CNVs were predominantly called in unique genome locations (94% for HapCNV and 97% for LUMPY, **Fig. S11A**), supporting their novel nature. This pattern was consistent when we randomly downsampled all sequencing data to the lowest read coverage prior to CNV calling (1.3 million reads, **Fig. S11B**). This comparison not only highlights the suitability of the common and rare categories but also the difference between the CNV calling methods. Based on these observations, we assessed common and rare CNVs using both methods to capture the broadest view of stress effects on CNV generation.

### Genome-wide de novo CNVs increased following replication stress

When we compared de novo CNVs in genomes with and without replication stress, we found that results were consistent regardless of the CNV calling method (**Fig. 2**). While the proportion of duplication and deletions somewhat changed with treatment (p value of 0.02 for decreased duplications in treated samples for LUMPY, **Fig. 2C**), we identified large differences in numbers de novo CNVs between treated and untreated 2-cell samples (p value of 0.0002 for HapCNV and <0.0001 for LUMPY, **Fig. 2D**). This pattern was consistent when we downsampled all sequencing data (p value of 0.001 for HapCNV and <0.0001 for LUMPY, **Fig. S12A & B**), indicating that the difference in de novo CNV counts between treatments was not due to read coverage. When we assessed common and rare CNV categories, we once again observed highly significant differences between treated and untreated 2-cell samples in common CNVs using both methods (common: p value 0.0004 for HapCNV and <0.0001 for LUMPY, **Fig. 2E**; rare: p value of 0.008 for HapCNV and <0.0001 for LUMPY, **Fig. 2F**).

When we compared the proportion of de novo CNVs relative to total CNVs per sample, rare CNVs were significantly increased over common CNVs (p value of 0.003 for HapCNV and 0.009 for LUMPY, **Fig. 2G**). This difference persisted regardless of downsampling (p value of 0.02 for HapCNV and 0.004 for LUMPY, **Fig. S12B & C**) and is in line with our assessment above that rare CNVs sit in unique genome locations and are more likely to be novel in nature (**Fig. S11**). Overall, we detected a ∼2-3-fold increase of de novo CNVs in treated samples, regardless of the CNV calling method (**Table 3**). Once again, rare CNVs displayed the largest increase following treatment (∼3-4-fold, **Table 3**, **Fig. 2H**).

### High-confidence de novo CNVs across the genome share consistent breakpoint characteristics

When we compared overlaps between the HapCNV and LUMPY (**Fig. 3A**), we detected a set of CNV regions that was consistent within sample groups (5 for untreated and 38 for treated, **Table S5**). The frequency of these “high-confidence” CNV regions also displayed an increase following replication stress (with knowns excluded, ∼12-fold increase in treated samples, **Fig. 3B**). Overall, high-confidence CNV regions represented both duplications and deletions (**Fig. 3B**) and were located on the majority of chromosomes (12 of 14, **Fig. 3C**). Of note, approximately half of these regions were identified as “rare” by both CNV calling methods across treated samples (17/36, 47%; **Table S5**), indicating that novel CNVs were stimulated in parasite genomes under stress.

**Figure 3:**
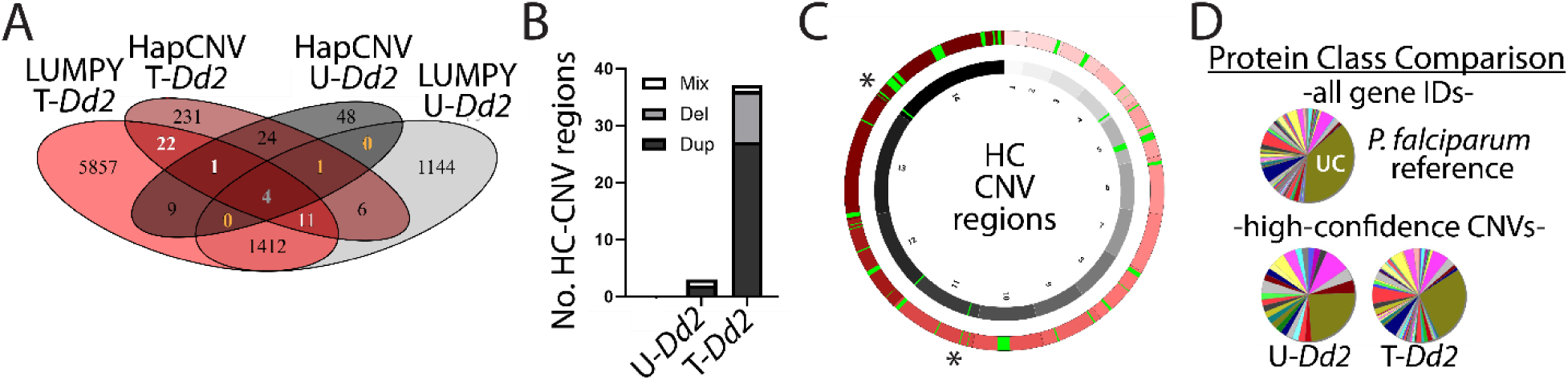
High-confidence CNVs are located across the genome and represent diverse protein classes. **A**. Comparison of CNV calls showing the number of CNV regions consistent across the two CNV calling methods. High-confidence CNV regions in untreated samples (U-*Dd2*, in yellow text); high-confidence CNV regions in treated samples (T-*Dd2*, in white text). Central number (grey): CNV regions consistent across all samples and calling methods (includes 2 known CNVs and 2 de novo CNVs, **Table S5**). **B**. Summary of number of high-confidence (HC) CNV regions in untreated (U-*Dd2*) and treated (T-*Dd2*) samples including duplication (Dup), deletion (Del), and mixed calls (sub-regions were called as duplications and deletions across a single CNV region by the same CNV calling method, i.e. HapCNV or LUMPY). **C**. Chromosomal location of high-confidence CNV regions identified by both HapCNV and LUMPY methods (green) from untreated (black) and treated (red) parasites. Only core regions of the genome are included in the representation; subtelomeric regions as defined by Otto, et al. were omitted. Each CNV region was increased in length by a factor of 2 to facilitate visualization relative to the rest of the genome. *, de novo high-confidence CNV regions identified in both untreated and treated samples. **D**. Panther classification system v19 protein class comparison. Top chart: protein classes from all annotated *P. falciparum* genes. Bottom charts: protein classes represented by high-confidence CNV regions in untreated (U-*Dd2*) and treated (T-*Dd2*) samples. UC: unclassified proteins (green). Other colors are randomly assigned by the program to represent diverse protein classes.

We also identified the proposed boundaries for the start and end of high-confidence CNV regions (termed “breakpoints”) and assessed their properties. Although CNV calling methods vary in their ability to precisely call CNV breakpoints (93), we were interested in whether these sequences shared characteristics with *P. falciparum* those identified using conventional methods (i.e. high AT content and nearby stable DNA secondary structures, (40)).). Overall, we found that features of breakpoints from high-confidence CNVs (12 for untreated and 66 for treated conditions, **Table S6**) were consistent with what was previously reported. Most breakpoints (>75%) had nearby stable secondary structures predicted to form hairpins with an average free energy of folding of −7 to −8 kcal/mol (depending on the treatment, **Table S6**). Most of these regions were situated in intergenic areas of the genome (56% for untreated and 75% for treated). The breakpoint regions also exhibited a higher AT-content compared to the genomic average (85-86%, **Table S6**, versus 80.6% genome-wide (41)). Based on breakpoint positions, we estimated the size of the high-confidence CNV regions; in general, this list represented relatively large regions (>4kb) that had the potential to cover multiple genes (maximum of 115kb, average of 23kb, **Table S6**).

#### High-confidence de novo CNVs represented diverse cellular pathways with potential clinical benefits

When we searched for genes that are covered by these regions, we identified 26 genes (across 3 de novo CNV regions) and 198 genes (across 37 de novo CNV regions) in untreated and treated *Dd2* samples, respectively (**Table S5**). Genes encompassed by the CNV regions represented diverse protein classes (**Fig. 3D**) and no gene ontology (GO) categories were significantly enriched in this list of genes (using an FDR adjusted p value of 0.05, **Table S7**). Along with the distribution across the genome (**Fig. 3C**), the lack of GO term enrichment emphasizes the random nature of the de novo CNVs and the absence of selection during sample preparation.

High-confidence CNV regions encompassed genes that were previously reported as important for clinical malaria infections (**Table 4**). Several genes play essential roles in transmission of the parasite from the human blood to the mosquito and back. We identified CNV regions that included genes for two TRAP-related proteins, CTRP and S6, which are important for infection of the mosquito midgut and salivary glands, respectively (94,95). Additionally, CNV regions encompassed two PHIST genes (Pfg14-744 and 748) that are important for early gametocyte development (96,97). We detected CNV regions covering 3 genes important for liver stage development including two genetically-attenuated vaccine candidates, UIS3 and SLARP (98–101); the latter of which achieved equivalent protection levels to whole sporozoite vaccines (PfSPZ-GA1, (102,103)). Two CNV regions carried genes important for replication of the erythrocytic stage (SEA1 and eif4A) and another two for virulence and antigen expression (PTP6 and SURFIN 14.1). One CNV region encompassed an aminophospholipid transporter previously identified as important for artemisinin resistance in a GWAS study (104). Finally, some of the genes from high confidence CNV regions were shown to be essential in the erythrocytic stage transposon mutagenesis screen (Pfg14-744, Pfg14-748, eif4A, and PTP6), indicating their multiple important roles across the parasite developmental cycle.

**Table 4.**
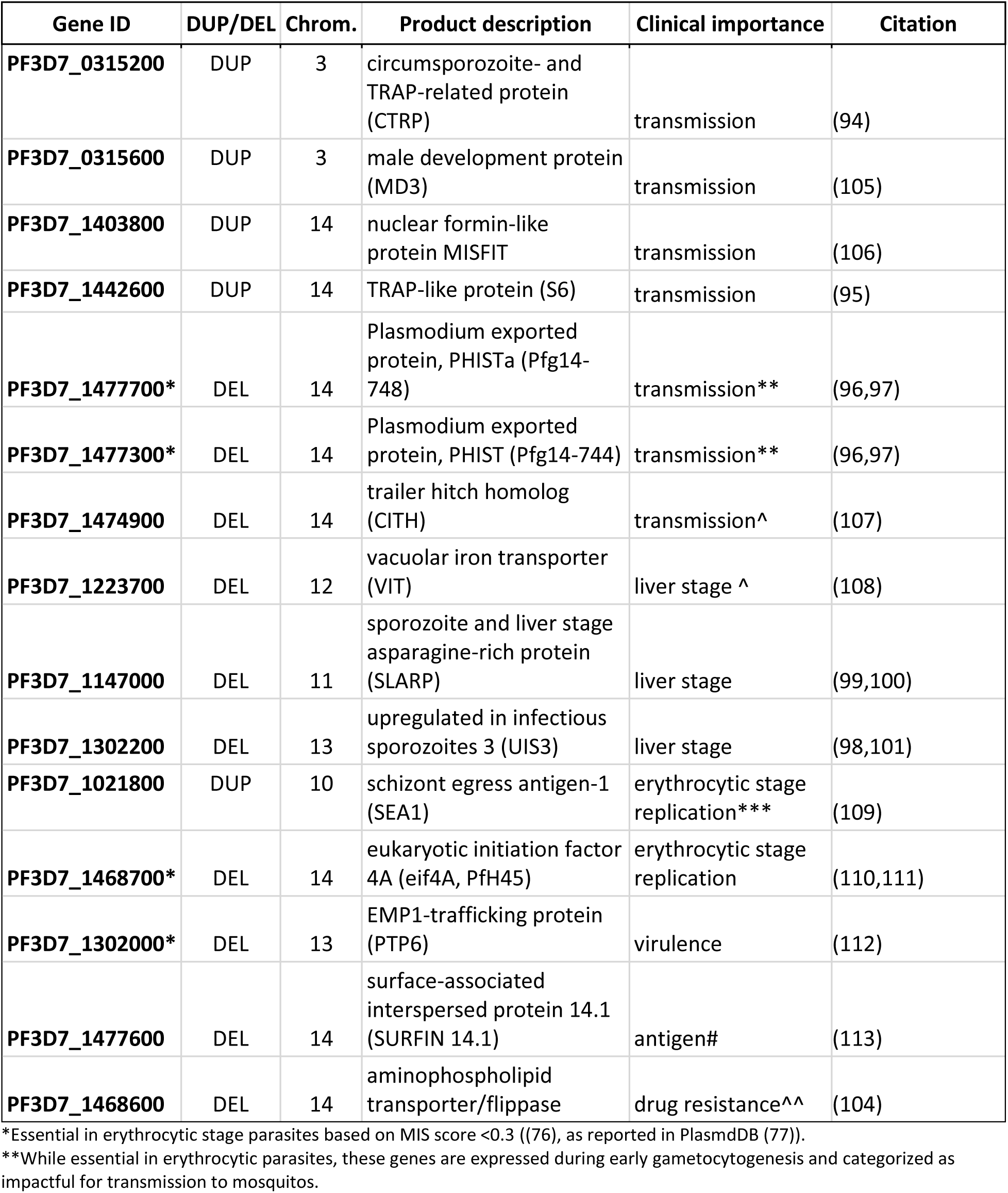
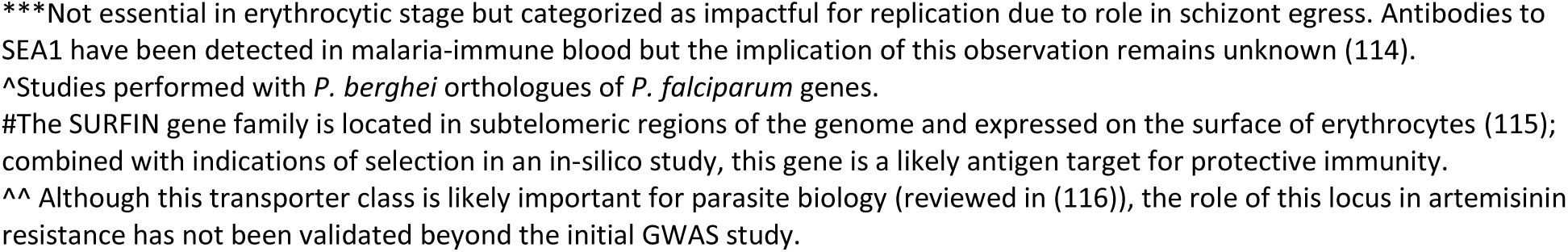
High-confidence CNV regions with clinical importance.

| Gene ID | DUP/DEL | Chrom. | Product description | Clinical importance | Citation |
| --- | --- | --- | --- | --- | --- |
| PF3D7_0315200 | DUP | 3 | circumsporozoite- and TRAP-related protein (CTRP) | transmission | (94) |
| PF3D7_0315600 | DUP | 3 | male development protein (MD3) | transmission | (105) |
| PF3D7_1403800 | DUP | 14 | nuclear formin-like protein MISFIT | transmission | (106) |
| PF3D7_1442600 | DUP | 14 | TRAP-like protein (S6) | transmission | (95) |
| PF3D7_1477700* | DEL | 14 | Plasmodium exported protein, PHISTa (Pfg14-748) | transmission** | (96,97) |
| PF3D7_1477300* | DEL | 14 | Plasmodium exported protein, PHIST (Pfg14-744) | transmission** | (96,97) |
| PF3D7_1474900 | DEL | 14 | trailer hitch homolog (CITH) | transmission^ | (107) |
| PF3D7_1223700 | DEL | 12 | vacuolar iron transporter (VIT) | liver stage ^ | (108) |
| PF3D7_1147000 | DEL | 11 | sporozoite and liver stage asparagine-rich protein (SLARP) | liver stage | (99,100) |
| PF3D7_1302200 | DEL | 13 | upregulated in infectious sporozoites 3 (UIS3) | liver stage | (98,101) |
| PF3D7_1021800 | DUP | 10 | schizont egress antigen-1 (SEA1) | erythrocytic stage replication*** | (109) |
| PF3D7_1468700* | DEL | 14 | eukaryotic initiation factor 4A (eif4A, PfH45) | erythrocytic stage replication | (110,111) |
| PF3D7_1302000* | DEL | 13 | EMP1-trafficking protein (PTP6) | virulence | (112) |
| PF3D7_1477600 | DEL | 14 | surface-associated interspersed protein 14.1 (SURFIN 14.1) | antigen# | (113) |
| PF3D7_1468600 | DEL | 14 | aminophospholipid transporter/flippase | drug resistance^^ | (104) |

\*Essential in erythrocytic stage parasites based on MIS score <0.3 ((76), as reported in PlasmdDB (77)).
\*\*While essential in erythrocytic parasites, these genes are expressed during early gametocytogenesis and categorized as impactful for transmission to mosquitos.
\*\*\*Not essential in erythrocytic stage but categorized as impactful for replication due to role in schizont egress. Antibodies to SEA1 have been detected in malaria-immune blood but the implication of this observation remains unknown (114). ^Studies performed with *P. berghei* orthologues of *P. falciparum* genes.

As a final analysis, we evaluated whether genes from high-confidence CNV regions specifically from our treated samples have been detected in parasites from clinical infections. We performed this analysis by searching the largest catalogue of clinically relevant CNVs to date; this list of high-frequency CNVs was previously called using genomes from 2855 parasite isolates from 21 malaria-endemic countries and represented genes from larger (>300bp), high-quality, core genome variants (38). Assuming ∼5000 genes in the core *P. falciparum* genome (58), we found that genes from the two lists were ∼2-times more likely to overlap than by chance (chi-square odds ratio of 2.5, Fisher’s exact p value <0.0001 for HapCNV-treated CNV list and 1.6, 0.03 for LUMPY-treated CNV list). Therefore, stress-induced de novo CNVs have the potential to be beneficial in the clinical environment.

## DISCUSSION

### Low-input genomics for studying de novo CNVs

Non-parental or “de novo” CNVs are not detected when analyzing a population of parasites predominantly because their signal (e.g. extra reads that align to that region or reads that span breakpoints) is negated by the overwhelming signal from normal copy number at that genome location. For this reason, assessments of fewer cells are necessary to investigate de novo CNV generation. While de novo CNVs have been previously tracked using flow cytometry of single yeast genomes (23), this sensitive approach provides a limited view of CNV dynamics by focusing on few specific loci that express fluorescent reporters. Inferring gene copy number through single cell transcriptomics can identify Mb-sized structural changes across genomes from heterogeneous tumor samples (117–120) but this approach is not applicable for smaller sized de novo CNVs. Single-cell genomics, where individual genomes are isolated and amplified to a level that can be sequenced, has been used to directly quantify de novo CNV rates in brain tissue, cancer cells, and *Leishmania* parasites (121–128). Early single-cell genomics studies of *P. falciparum* have been promising but so far have had limited success with CNV identification (42,129,130). Here, we optimized a low-input analysis pipeline and successfully isolated, amplified, and sequenced *P. falciparum* samples for CNV analysis. With experimental and computational improvements, we were able to increase our rate of parental, or “known”, CNV calling over our prior study (42). Importantly, we detected de novo CNVs across the parasite genome and replication stress significantly increased their rate of formation. By analyzing ∼45 low-input samples per condition, we are limited to observing a subset of the population. However, this study is one of few that have directly assessed de novo CNVs in microbes and our findings demonstrate that replication stress readily drives the rapid generation of de novo CNVs in an important pathogen. Below, we cover the rationale behind our experimental/analysis choices and integrate our findings into an overarching model of *P. falciparum* adaptation.

### Application of stress without evidence of selection to explore CNV dynamics

To evaluate the impact of replication stress on de novo CNV generation, we applied treatment to parasites just prior to replication. We observed replication stall and then resume post-treatment, which provided evidence that we successfully applied sub-lethal stress (**Figs. 1D**, **S1**, and **S2**). Following this step, we allowed the parasites to complete replication and reinvade new erythrocytes; we reasoned that this “recovery phase” enabled the repair of the resulting DNA damage, which is likely to be replication-dependent (reviewed in (1,131). Additionally, reinvasion facilitated the isolation of haploid parasite genomes (1n, **Fig. 1A**), encouraging the detection of de novo CNVs due to limited contrasting signal (26).

Because of the reinvasion step, which involves an expansion in parasite number (∼3-fold, **Fig. 1D**), there was potential to select for beneficial DNA changes across the population of parasites. However, we did not detect evidence of strong selection from SNP profiles (**Fig. S10A**) or high-confidence CNV regions (**Fig. 3**, **Table S5**). Specifically, we did not observe an enrichment of CNVs that encompassed the *pfdhodh* gene, which contributes directly to DSM1 resistance (132), or genes from DNA-related pathways (**Table S7**). While studies of numerous microbes observe CNVs arise under strong selective conditions like nutrient starvation or drug treatment (as for *Saccharomyces* yeast: (5,133,134), *Salmonella* bacteria: reviewed in (135), as well as *Plasmodium*: (132,136–142)), we now show that mild, transient conditions can stimulate CNVs that have the potential to increase parasite survival.

### Relative comparison using multiple CNV calling methods to appreciate the impact of stress

De novo CNV estimates using single cell methods from yeast, neurons, and human cancer cells vary greatly and are difficult to standardize due to the use of different experimental techniques and CNV calling methods (23,24,143). For these reasons, we are not attempting to compare the rate of *P. falciparum* CNV formation from this study to those from other organisms. Additionally, this lack of standardization in the field led us to use two distinct CNV calling methods in our analysis. Due to the strengths and weaknesses of HapCNV and LUMPY, we observed differences in both known (**Fig. 2B**) and de novo CNV (**Table S4**) calling using the two methods. LUMPY identifies reads that cover breakpoint regions to sensitively detect CNVs (70); because we are counting regions with few reads as support, both sensitivity and the number of false positives are high in this analysis. High known calling rates along with high numbers of de novo CNVs in our studies exemplified this feature of LUMPY. On the other hand, HapCNV uses a genome-specific pseudo-reference for normalization, which removes repeated patterns of over-and under-amplification ((44) and *Materials and Methods*); because we require read coverage to span 3 consecutive 1kb bins, small CNVs in lower coverage genomic neighborhoods are excluded in HapCNV analysis. This limited the detection of smaller known amplicons (*pf11-1* and *pf332*), led to fewer de novo CNV calls using this method, and likely contributed to the larger size of high-confidence CNVs (**Table S6**). Given the high abundance of small CNVs (<300bp) in the parasite genome (38,144), HapCNV is likely underestimating the impact of small CNVs in our study.

Ultimately, the value of our study is in the relative comparison of treated and untreated samples. During sample preparation, we acknowledge that the apparent 3-fold difference in amplification efficiencies between untreated and treated sample groups (**Fig. S6**) could compromise this comparison. In order to prevent cross-contamination, 2-cell samples were sorted into different plates and consequently, they were amplified separately. Although separated by just a few days (see *Materials and Methods*), alterations in reagent performance from batch to batch could contribute to differences in amplification efficiency. Additionally, we speculate that there may have been a general under-quantification of the treated *Dd2* plate; we increased the amount of each sample sequenced by 4-fold to compensate for low DNA amounts and ended up achieving ∼4-fold higher coverage levels for treated samples (**Table 2**). This under-quantification is supported by our ddPCR assays, where samples with small amounts of quantified DNA still yielded abundant PCR positive droplets (**Fig. S7D**), indicating adequate *P. falciparum* genome amplification for this round. Because sequencing quality was similar between sample groups (**Table 2**, CV and mapping quality), downsampled data showed a consistent result with analyses using all data (**Figs. 2**, **S11**, and **S12**), and treatment had a consistent impact on CNV categories using two distinct CNV calling programs (**Table 3** & **Fig. S12B**), we are confident that coverage differences are not dramatically impacting our overall conclusions.

During our analysis, we assumed that false positive CNVs occur at a similar rate across both treated and untreated groups (within each CNV caller and coverage level), which allowed us to confidently assess the impact of stress despite some experimental variation. Variations in known and de novo CNV calling described above emphasize that no CNV calling method is perfect and combining them can improve confidence in results (42,145–147). Therefore, we investigated de novo CNV patterns using the two individual methods (**Figs. 2**, **S11**, & **S12**) as well as those that overlapped between HapCNV and LUMPY (**Fig. 3**, **Table S5**). Importantly, these “high-confidence” CNVs reflected increases after stress previously detected using the individual tools, albeit at a greater level (∼3-versus 12-fold increase). Additionally, we speculate that newly arising CNVs would have distinct locations across samples; thus, the rare nature and unique locations of high-confidence CNVs emphasized their potential to be novel (**Table S5**).

### De novo CNV categories highlighting existing and novel genome variation

During our investigations, we identified two types of de novo CNVs; those detected in one or a few genomes (rare, <10%) and those detected in more than a few (common, >10%). While others have used these CNV categories (73), there is no precedence for them in the context of *Plasmodium* biology (i.e. a haploid parasite with asynchronous replication and schizogeny (148)). However, we propose that tracking these categories helps us to understand the biological relevance of de novo CNVs in our analysis.

Based on their frequency, common CNVs are either artifacts of low-input procedures/CNV analysis or represent minor variants that preexisted in the population or arose early in the replication cycle. For the former, bias during the whole genome amplification step (i.e. the repeated pattern of over/under-amplification that occurs in a reproducible pattern across the parasite genome (42)) and PCR during library construction have the potential to skew gene copy number and increase the false positive rate (149,150). However, we chose experimental and computational methods designed to limit the contribution of amplification bias. First, MALBAC amplification itself limits the over-amplification of certain genomic regions by avoiding exponential amplification at the earliest steps (149) and we used limited PCR cycles during library preparation (3 cycles, (42)). These efforts are most clearly shown through the reduction in CV following MALBAC optimization in both of our studies (by ∼39% after modifying the amplification primer (42) and by 43% after switching to the *Bsu* polymerase, **Table S1**). Second, LUMPY is not dependent on read coverage and HapCNV specifically addresses amplification artifacts by removing repeated signal present in all samples (44,70). Overall, we detected few CNVs with conserved genomic locations across low-input samples, which provides evidence that our methods limit the effect of amplification bias on the results; we only identified two high-confidence CNV regions that had conserved locations across multiple *Dd2* and *FCR3* 2-cell samples (**Fig. 3C**, **Table S5**). In the future, amplification-free methods (151), or visualization of single long-reads (49,152), may offer advantages in distinguishing amplification bias from minor variants and de novo CNVs.

Rare CNVs, on the other hand, represent either random noise or true signal from novel CNVs arising in the genome. We assert that most noise is removed through normalization procedures, especially with HapCNV, and the impact of remaining false positives are minimized by the relative comparison of our studies (see above). We identified the majority of rare CNVs in unique genome locations across sample types, providing evidence that they are not a result of amplification bias where the same CNVs are repeatedly detected in each sample. Additionally, the greater impact of stress on rare CNVs than common CNVs (**Table 3** and **Figs. 2F, 2G**) supports their replication dependence. The random nature of de novo CNVs, as well as the capacity to encompass any gene across the genome, ensures that CNVs can alter all aspects of parasite biology in response to the host environment. Further, our finding that stress-induced de novo CNVs tended to exhibit altered copy number in clinical isolates combined with the high frequency of unique CNVs in previous genome-wide CNV studies (32,38), directly illustrate the expansive evolutionary potential of this pathogen.

### Adaptations that encourage de novo CNV formation

The current model of CNV formation in asexual erythrocytic *P. falciparum* parasites is that AT-rich sequences form hairpins, disrupt replication, and eventually lead to double-strand breaks that are repaired by error-prone pathways (40). The evolution of CNVs in this organism is especially interesting because of its unique genome architecture and alternative repertoire of CNV-generating repair pathways (41,153). Although they arise at many locations across the genome ((30,31,38), **Fig. 3C**), *P. falciparum* CNVs that contribute to adaptation are commonly gene duplications with a relatively simple structure. Many impactful duplications form in tandem head-to-tail orientation ((40,132,142,154), **Fig. 4A**), which is likely due to a limited repertoire of DNA repair pathways; *P. falciparum* lacks the canonical nonhomologous end-joining (NHEJ) pathway that is a major contributor to CNV formation in other organisms (1,153). Instead, parasites use pathways that employ varying lengths of sequencing homology (i.e. homologous recombination, or HR, and microhomology-mediated repair, **Fig. 4B**). This repair repertoire, along with an especially high AT-content genome that facilitates CNV formation (40,132), and a lack of cell cycle checkpoints that control replication forks during times of stress (reviewed in (155)), likely represent adaptations that assist haploid *P. falciparum* parasites in accumulating CNVs across their genome (**Fig. 4A**). One major question that remains is whether these same processes are active in other rapidly replicating parasite stages such as oocysts in the mosquito midgut or schizonts in the human liver.

**Figure 4:**
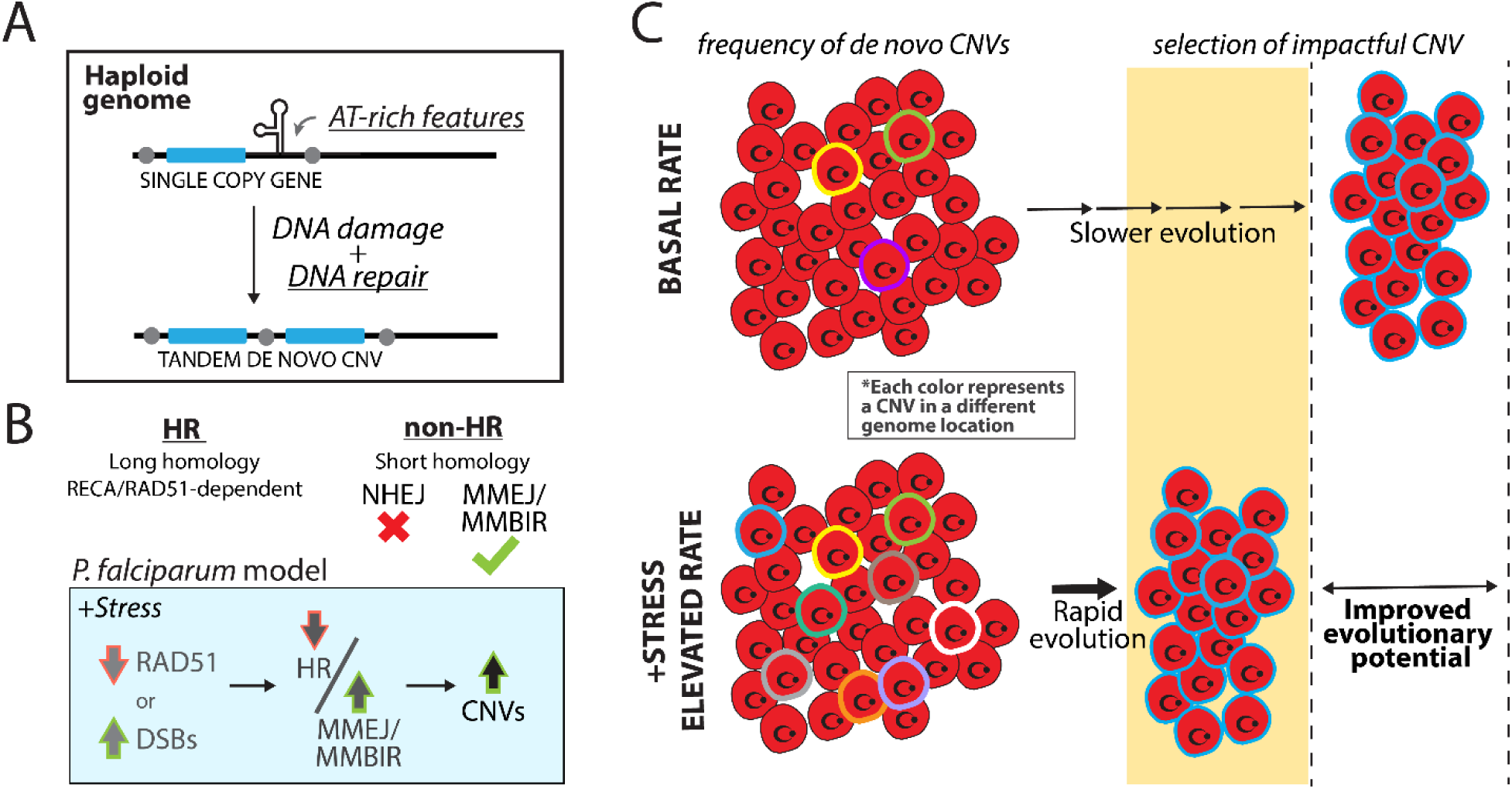
**Potential connection between replication stress, DNA repair, and CNV generation in the malaria genome**. **A**. Adaptations that encourage CNV formation in the *P. falciparum* genome (underlined, (40)). **B**. Proposed model of how replication stress can impact DNA repair pathways based on prior studies (HR, homologous recombination; NHEJ, non-homologous end-joining; MMEJ, microhomology-mediated end joining; MMBIR, microhomology-mediated break-induced repair; DSB, double-strand breaks). Model is suggested based on studies showing 1) stress can lead to reduction of RAD51 and increased DSBs reduced HR activity (156,157), 2) stress can trigger pathways that use microhomology sequences for DNA repair (158,159), and 3) *P. falciparum* uses microhomologous sequences to generate CNVs (40). **C.** Potential benefits of a diverse parasite population for evolutionary potential. Stress elevates the frequency of de novo CNVs across the population, which leads to more rapid evolution of beneficial CNVs (blue cells).

### Updating the model of P. falciparum genome adaptation

In conjunction with previous studies in other organisms, our findings in *P. falciparum* support the connection between replication stress, DNA repair, and CNV generation (**Fig. 4B**). Studies from bacteria to cancer cells have shown that stress can either alter levels of proteins essential for HR-based repair or increase the frequency of DNA breaks (159). In cancer cells, hypoxic stress causes a decrease in HR activity (156) and this may lead to an increased reliance on alternative error-prone pathways to repair DNA damage (160). In bacteria (158,161), starvation drives initial amplification steps at microhomologous sequences. The nature, degree, and timing of applied stress likely matter because some studies also show the dose-dependent induction of HR proteins under some conditions (162). Studying these processes in *P. falciparum* under various conditions will be particularly important due to the lack of NHEJ repair (see *Adaptations that encourage de novo CNV formation*); when NHEJ is deficient in cancer cells, DNA repair becomes more error-prone and contributes to CNV formation (163,164).

Prior to the current study, the predominant evidence connecting DNA repair and CNV generation in *P. falciparum* was the detection of microhomology-mediated pathway signatures in CNV breakpoints (40). Microhomology-mediated pathways require less homology and therefore, are more likely to interact with diverse sequences up- and downstream of a DNA break to generate various length CNVs. Our current observation of stress-induced de novo CNV formation (**Table 3** and **Fig. 2H**) further supports this model. We originally set out to understand how sub-lethal antimalarial treatment impacts CNV generation because this is a condition that the parasite encounters during clinical infection (165). However, because this compound targets pyrimidine biosynthesis and its impacts mirror aphidicolin treatment (**Fig. S1**), we speculate that the effects we observed are likely due to replication inhibition. Our result is consistent with studies on diverse organisms, like humans, mice, and flies, where replication stress leads to CNV formation (20–22,166,167). Interestingly, the degree of de novo CNV stimulation is also consistent across these studies; we detected a ∼3-fold increase in *P. falciparum* de novo CNVs (**Table 3**) compared to a ∼3-5-fold increase in mammalian de novo CNVs (20–22). Compared to this work, our study highlights that despite the smaller genome and divergent biology of *Plasmodium*, as well as the use of different experimental approaches, there are likely conserved processes that increase genetic diversity under stress. Future studies will address gaps in knowledge including assessment of conditions that stimulate the *P. falciparum* adaptive amplification response, the timing of this response, and the requirement and regulation of specific mechanistic players.

### Support for adaptive amplification in microbial populations

The experimental evolution approach to evaluate the impact of selective conditions on microbial populations across relatively long time scales has provided many insights that include the important role of gene amplification in genomic diversity (e.g. (5,134)). In bacteria, the term “adaptive amplification” was established to describe how environmental conditions can induce genetic change to help cells adapt to stress (161). De novo CNVs have been directly investigated in bacteria and *Leishmania* parasites using single read assessments and single-cell isolation, respectively (128,152); to our knowledge, prior studies have not assessed the effect of non-selective short-term conditions on de novo CNV rates in microbes. By continuing to improve genome amplification and CNV detection methods, we anticipate that future studies of genome dynamics will lead to new insight on how stress stimulates microbial evolution.

Already, studies from a variety or organisms are showing that an increased rate of CNV formation can have an impact on a population of organisms; if beneficial under specific conditions, a CNV that arises in a single genome can expand during selection into a larger population of cells with novel characteristics (135,168). This rapid expansion is exemplified when minor bacterial populations with higher gene copy numbers confer “heteroresistance” during clinical antibiotic selection (152,169–171). In another example, higher levels of intra-tumor heterogeneity in gene copy number predict a poorer cancer prognosis (127,172,173).

For *Plasmodium*, even with a change in the copy number of a single region per parasite, the genomic diversity within a single infected human is expansive due to the sheer numbers of parasites (estimated to reach 10^8^ parasites when symptomatic and >10^11^ in severe *P. falciparum* infection (174)). This diversity can provide an obvious advantage as a heterogeneous population prepares asexual parasites to respond to diverse stressors (**Fig. 4C**). While our current studies point to a role of adaptive amplification in *P. falciparum*, we do note a low level of de novo CNVs across the parasite genome under normal conditions (**Fig. 2D** and **S12A**). Since some antimalarials act rapidly (81), some level of de novo CNVs already present in a few parasites across the population would increase the chances of survival during drug treatment. It is also important to understand how parasites respond to stressful environments during infection, including changes in nutrient composition in different hosts, drug treatment during symptomatic infection, or attack from the human immune system. While the current study focused on one antimalarial compound, it will be important for future studies to evaluate the impact of other sources of stress on *P. falciparum* CNV formation. For example, hypoxia stimulates CNV formation in cancer cells (156) and a proteotoxic drug stimulates genetic change in yeast (175). As a eukaryotic microbe, *P. falciparum* occupies special niche that can be exploited as a model to study eukaryotic mechanisms of stress-induced genome dynamics.

### Clinical implications & future questions

*P. falciparum* causes the majority of malaria deaths worldwide and readily acquires antimalarial resistance (176,177). Resistance-conferring CNVs that encompass multiple genes have been identified in clinical infections (30,38,178–182). While a gene from one of our high-confidence CNVs may contribute to artemisinin resistance, we also identified genes that participate in many processes important for malaria transmission and infection (**Table 4**). This observation combined with the diverse genome location and protein classes represented (**Fig. 3**) illustrates the enormous adaptive potential of the *P. falciparum* genome.

Despite their direct contributions to various phenotypes, CNVs may also facilitate the acquisition of point mutations in haploid *P. falciparum*; strong evidence for the close relationship comes from the observation of point mutations *within* amplifications selected in vitro (137,139,142,183,184). Once de novo CNVs form during replication of the asexual erythrocytic stage (**Fig. 4**), meiotic recombination during the sexual phase in the mosquito can streamline beneficial CNVs to balance fitness costs (68). Given the importance of CNVs in *P. falciparum* adaptation, it is not surprising that this organism has evolved strategies to encourage CNV formation (as described in *Adaptations that encourage de novo CNV formation*). Additionally, parasites from specific regions of the world may have an increased propensity to develop drug resistance (185). Evaluating whether the CNV rate correlates with the parasite background will help to define the evolutionary potential of this successful pathogen.

Antimalarial therapies and vaccines targeting the *Plasmodium* parasite are in danger due to drug resistance and access challenges (186,187). A strategy to impede genome evolution may be required to control malaria infections. “Evolution-proof” therapies have been explored in response to antibiotic and anticancer resistance (reviewed in (188,189)). Of particular note is a recent study targeting NHEJ repair to prevent resistance in melanoma cell lines (190). While antimalarial combination therapies were originally suggested to limit recrudescence following artemisinin-based drug treatment (191), the parasite has gained CNV-based strategies to overcome partner drugs ((181,182) and reviewed in (192)). Therefore, identifying and targeting unique aspects of *Plasmodium* biology that facilitate CNV formation (e.g. DNA repair, AT-dependent mechanisms, or replication control) may limit parasite evolution and increase treatment efficacy.

## Data Availability

The source code for the BLAST pipeline and SVCROWS can be accessed at https://doi.org/10.6084/m9.figshare.29885879 and https://doi.org/10.6084/m9.figshare.29099801.v1. Short read data is available at NCBI Sequence Reads Archives under project PRJNA1201106.

## Supplementary Data are available at *NAR* Online

## Acknowledgments

We would like to thank Dr. John Campbell for the use of the Mosquito LV instrument and assay reagents for the colorimetric assessment of the SH800. We would also like to thank Dr. Ali Guler for statistical consultation and Dr. Heidi Seears from the University of Virginia Department of Biology Genomics Core Facility for sequencing assistance. We also thank the Vector and Eukaryotic Pathogen Genomics Database Resource (VEuPathDB) and specifically PlasmoDB (https://plasmodb.org) for hosting data used in some aspects of this study.

