## Supplementary material for "Replication stress increases de novo CNVs across the malaria parasite genome": Fig. S1-12, Table S1

### SUPPLEMENTAL FILES

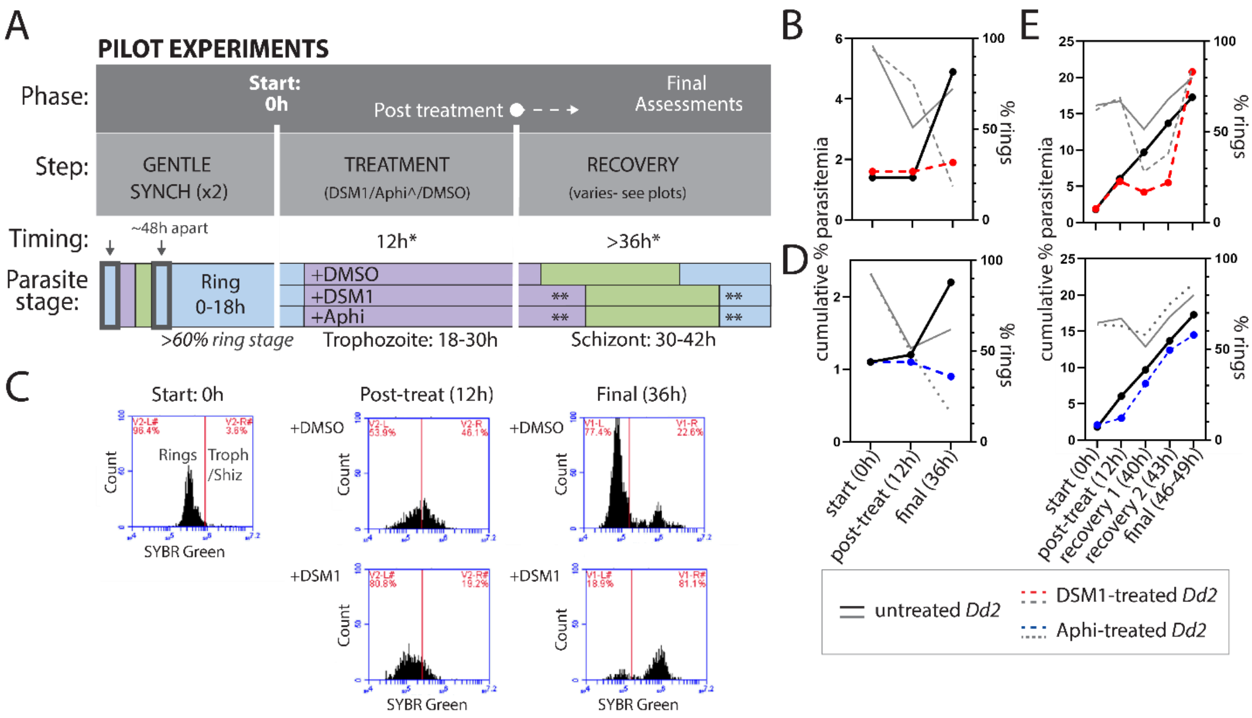


**Figure S1: DSM1 reversibly inhibits parasite growth during pilot short-term sublethal treatments. A**. Timeline of parasite treatment and recovery for independent pilot experiments assessing DSM1 and aphidicolin treatment compared to solvent control (dimethylsulfoxide, DMSO). Parasites were synchronized to enrich for rings before short-term treatment was applied. Parasite growth and viability were tracked from start, to post-treatment, to final assessments. The recovery period allowed for completion of the erythrocytic cycle and reinvasion to assess growth rate and viability. *, cumulative time from start (0h). **, approximate delay in parasite staging following treatment with replication inhibitor. Aphi, aphidicolin. **B, D, E.** Cumulative parasitemia and ring percentages in pilot treatments of *Dd2* parasites with DSM1 (panels B, C, and E, top) and aphidicolin (panels D and E, bottom). Cumulative % parasitemia was calculated by multiplying the parasitemia by the last dilution factor for the period being measured (e.g. post-treatment). Ring % is calculated from flow cytometry plots tracking SYBR Green staining, where rings (1n) have lower florescence than late-stage parasites (>1n, see panel C). Timing is for each experiment is indicated on each plot. Standard error is calculated from duplicate or triplicate flasks but is too small to view on the plot (<0.33% in all cases). For all panels (see key): untreated *Dd2* (black/grey, solid lines), DSM1-treated *Dd2* parasites (red/grey, dotted lines), and aphidicolin-treated *Dd2* parasites (blue/gray, dotted lines). **B**. Effects of short-term DSM1 versus DMSO treatment, n=3. **C**. Flow cytometry histograms from representative samples from experiment in panel B showing stage progression of DMSO-(top) versus DSM1-treated (bottom) parasites using SYBR Green staining. Red line demarks boundary between ring and later stage parasites (trophozoites/schizonts); red number shows percentage of events that fall within the left (ring) or right (late-stage) gates. DSM1 treatment limits progression from ring to trophozoite stage (post-treat) and delays completion of cycle after recovery (final). Data shown from one representative flask each. **D.** Effects of short-term aphidicolin versus DMSO treatment, n=2. **E.** Effects of short-term DSM1 (top) and aphidicolin (bottom) versus DMSO treatment, n=2.

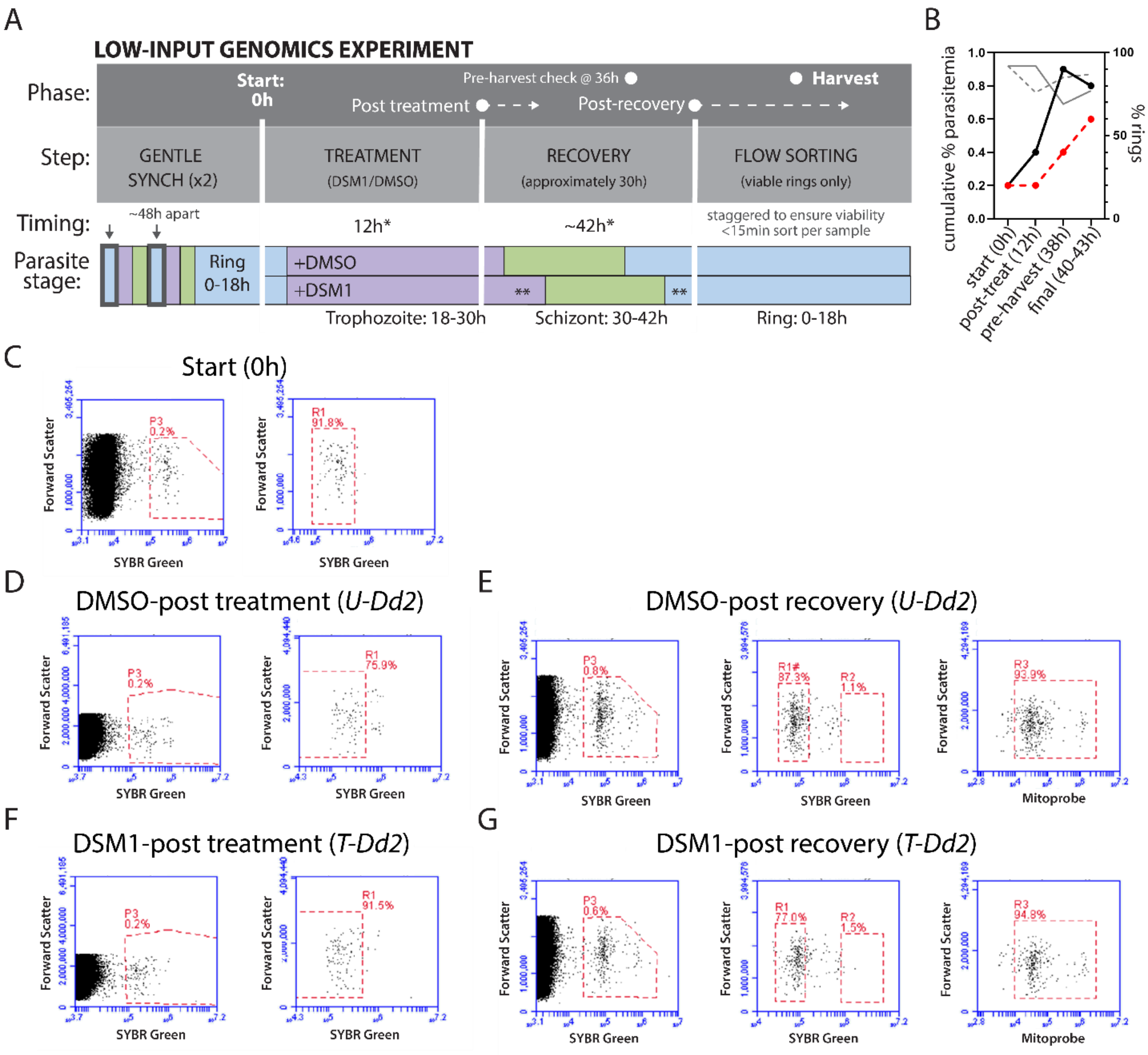


**Figure S2: DSM1 treatment for low-input genomics stalls parasite replication before recovery. A**. Timeline of parasite treatment and recovery for low-input genomics experiment. Parasites were synchronized to enrich for rings before short-term treatment was applied (DSM1 or solvent control, dimethylsulfoxide, DMSO). Parasite growth and viability were tracked from start, to post-treatment, to post-recovery, to harvest. The recovery period allowed for completion of the erythrocytic cycle and reinvasion to assess growth rate and viability. *, cumulative time from start (0h). **, approximate delay in parasite staging following treatment with replication inhibitor. **B**. Cumulative parasitemia and ring percentage in low-input genomics experiment. Cumulative % parasitemia was calculated by multiplying the parasitemia by the last dilution factor for the period being measured (e.g. post-treatment). Ring % is calculated from flow cytometry plots tracking SYBR Green staining, where rings (1n) have lower florescence than late-stage parasites (>1n, see **Fig. S1C**). Standard error is calculated from duplicate flasks but is too small to view on the plot. Untreated *Dd2* (black/grey, solid lines), DSM1-treated *Dd2* parasites (red/grey, dotted lines). **C-G.** Flow cytometry plots from representative *Dd2* samples: forward scatter (reflects cell size) vs SYBR Green (stains parasite genome) plots indicate the proportion of erythrocytes that contain parasites (% parasitemia, P3 gate) and the percentage that are ring (R1 gate) or late-stage parasites (R2 gate); forward scatter vs MitoProbe DiIc1 (5) (stains viable parasites) plots indicate the proportion of parasite containing erythrocytes (SYBR Green+) that are viable (R3 gate). Data shown from one representative flask each. **C**. Flow cytometry of *Dd2* parasites prior to treatment. **D**. Flow cytometry of *Dd2* parasites prior to treatment (start, 0h). **D**. Flow cytometry of untreated *Dd2* (*U-Dd2*) following incubation with solvent control (DMSO post-treatment). **E.** Flow cytometry of *U-Dd2* following ~30h of recovery (DMSO post-recovery). Recovery and reinvasion are evident from the increase in parasitemia, increase in ring stage, and high viability (>90%). **F**. Flow cytometry of treated *Dd2* (*T-Dd2*) following incubation with DSM1 (post-treatment). **G**. Flow cytometry of *T-Dd2* following ~30h of recovery (DSM1 post-recovery). Reinvasion is evident from the increase in parasitemia, increase in ring stage, and high viability (>90%).

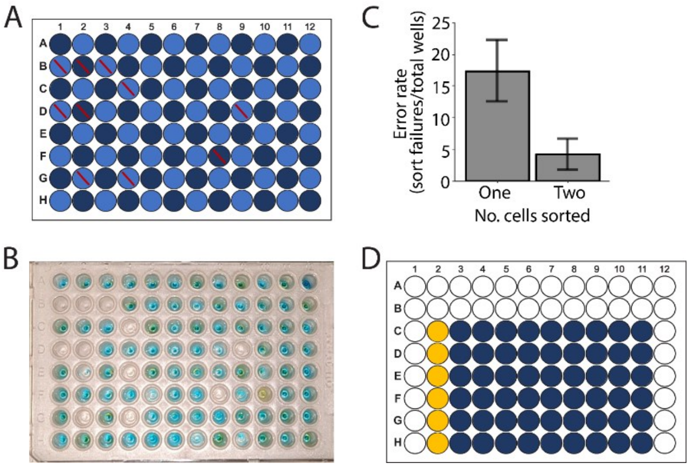


**Figure S3. Sorting low-input populations improves accuracy of parasite isolation. A**. FACS sorting plate diagram indicating how many cells were sorted per well and positions of failures. Dark blue: 2-cell sort, light blue: 1-cell sort, red slash: failed sort (see panel B for plate image). **B**. Representative sorting plate showing colorimetric results. Green, blue, and yellow color indicate positive sorting events. The plate had 2.5µl of reaction buffer prior to sorting to mimic low-input isolation conditions. **C**. Error rate for the number of cells sorted into each well. Error rate: sort failures (see panels A and B)/total wells of specific condition (i.e. 1- or 2-cells sorted per well). Error bars: SEM (n=3). **D**. Final sorting positions for low-input isolation. Top 2 rows and first and last columns of the plate were excluded. Orange: final 10-cell sort positions, blue: final 2-cell sort positions).

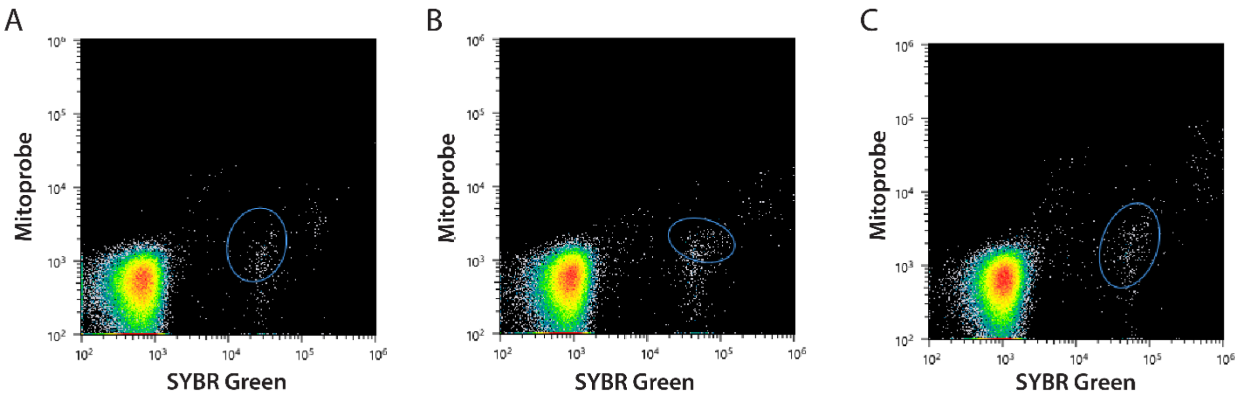


**Figure S4: FACS displays isolation of viable ring-stage parasites for low-input genomics.** FACS plots from the low-input genomics parasite isolation step (post-recovery, **Figs. 1C, S2A**); MitoProbe DiIc1 (5) (stains viable parasites) vs SYBR Green (stains parasite genome) plots indicate the proportion of erythrocytes (heat map of density) that contain viable ring stage parasites (blue circles were drawn after a test run of ~100 cells and reflect actual data points from sorting). All sample collections were staggered and limited to <15min to ensure parasite viability. Differences in ring-stage parasitemia from **Fig. S2E, G** are due to a slight delay in harvest times and a small drop in the numbers of viable parasites during collection. **A**. Untreated *FCR3* (blue circle: 0.27% ring stage parasitemia). **B**. Untreated *Dd2* (blue circle: 0.33% ring stage parasitemia). **C**. DSM1-treated *Dd2* (blue circle: 0.38% ring stage parasitemia, plot also depicted in **Fig. 1F**).

| *Pf*MALBAC Version 1 (2021) | *Pf*MALBAC Version 2 (current) |
| --- | --- |
| **parasite isolation** | |
| Microscopy based: Cell Microsystems CellRaft Air System | FACS-based: Sony SH800 Sorter |
| **MALBAC amplification** | |
| Primer: 20% GC random primer version 1: 5'GTGAGTGATGGTTGAGGTAGTGTGGAGNNNNNTTT 3' | Primer: 20% GC random primer version 2: 5'GTGAGTGATGGTTGAGGTAGTGTGGAGNNNNNNNNNNTTT 3' |
| DNA Polymerase: *Bst* large fragment (optimal at 60-65°C) | DNA Polymerases: *Bst* (60-65°C) & *Bsu* large fragment (optimal at 37°C) |
| Linear Cycling Parameters: 10°C – 45s, 15°C – 45s, 20°C – 45s, 30°C – 45s, 40°C – 45s, 50°C – 45s, 64°C – 10min, 95°C – 20s, 58°C– 1min between cycles | Linear Cycling Parameters: 10°C – 45s, 15°C – 45s, 20°C – 45s, 30°C – 45s, 64°C – 10min OR 37°C – 10min, 50°C 45s, 64°C – 45s, 95°C – 20s, 58°C– 1min between cycles |
| Pipetting: manual | Pipetting: robotic pipetting during linear steps (Mosquito LV) |
| Cycles: 19 linear, 17 exponential | Cycles: 2 linear (*Bst*), 19 linear (*Bsu*), 17 exponential |

**Figure S5: Changes in MALBAC amplification approach since previous publication.** *Pf*MALBAC version 1 details are as previously published as "optimized MALBAC" (42). *Pf*MALBAC version 2 details were employed in the current study to improve isolation and scale-up (microscopy vs FACS), limit contamination potential (manual vs robotic pipetting of additional enzyme for each round of linear cycling), and improve intergenic coverage (*Bst* vs *Bsu* DNA polymerase and changes in linear cycling parameters). Steps that are distinct in *Pf*MALBAC version 2 are underlined. FACS; fluorescence activated cell sorting.

A

| | 1 | 2 | 3 | 4 | 5 | 6 | 7 | 8 | 9 | 10 | 11 | 12 |
|---|---|---|---|---|---|---|---|---|---|---|---|---|
| A | | | | | | | | | | | | |
| B | | 4 | | | | | | | | | | |
| C | | 115 | 193 | 120 | 102 | 182 | nd | nd | nd | nd | nd | |
| D | | 68 | nd | 82 | nd | 122 | 142 | 120 | nd | nd | nd | |
| E | | 143 | 152 | nd | nd | 160 | 161 | nd | nd | nd | nd | |
| F | | 177 | nd | 147 | nd | nd | nd | nd | nd | nd | nd | |
| G | | 181 | nd | 114 | nd | nd | 216 | nd | nd | nd | 194 | |
| H | | 106 | nd | nd | 146 | 184 | nd | nd | nd | 174 | nd | |

B

| | 1 | 2 | 3 | 4 | 5 | 6 | 7 | 8 | 9 | 10 | 11 | 12 |
|---|---|---|---|---|---|---|---|---|---|---|---|---|
| A | | | | | | | | | | | | |
| B | | | | | | | | | | | | |
| C | | 214 | 137 | 67 | 148 | 152 | 174 | 161 | 400 | 165 | 138 | |
| D | | 135 | 138 | 132 | 134 | 129 | 109 | 112 | 130 | 160 | 117 | |
| E | | 178 | 172 | 154 | 144 | 130 | 161 | 124 | 194 | 136 | 149 | |
| F | | 131 | 180 | 170 | 170 | 183 | 120 | 130 | 142 | 187 | 176 | |
| G | | 199 | 126 | 127 | 170 | 142 | 124 | 144 | 136 | 136 | 130 | |
| H | | 180 | 125 | 149 | 146 | 150 | 113 | 272 | 133 | 146 | 106 | |

C

| | 1 | 2 | 3 | 4 | 5 | 6 | 7 | 8 | 9 | 10 | 11 | 12 |
|---|---|---|---|---|---|---|---|---|---|---|---|---|
| A | | too low | | | | | | | | | | |
| B | | | | | | | | | | | too low | |
| C | | 88 | 68 | 15 | 18 | 55 | too low | 10 | too low | 5 | 68 | |
| D | | 70 | 60 | 109 | 162 | 167 | too low | 74 | 74 | 10 | 8 | |
| E | | 8 | 5 | 140 | 71 | 139 | 100 | 3 | 13 | 47 | 46 | |
| F | | 86 | 50 | too low | 11 | 10 | too low | 25 | too low | 9 | 8 | |
| G | | too low | too low | 6 | 8 | too low | 67 | 44 | too low | too low | too low | |
| H | | 76 | too low | 7 | too low | too low | 22 | too low | too low | 79 | too low | |

**Figure S6: DNA quantity following MALBAC amplification is sufficient for sequencing (>10ng).** Plate maps show total DNA as measured by Qubit (total ng), Total DNA quantity indicated by color, green: >100ng, orange: 10-100ng, gray: <10ng or no cell added (column 1 and A/B2-12), nd: not determined. White text: Illumina sequenced, black text: not sequenced. Zero cells were sorted into the top two rows (A and B) and first column (1) of the plates to limit evaporation in low-input samples. 10-cell samples were confined to column 2. **A**. Untreated *FCR3* samples. **B**. Untreated *Dd2* samples. **C**. DSM1-treated *Dd2* samples.

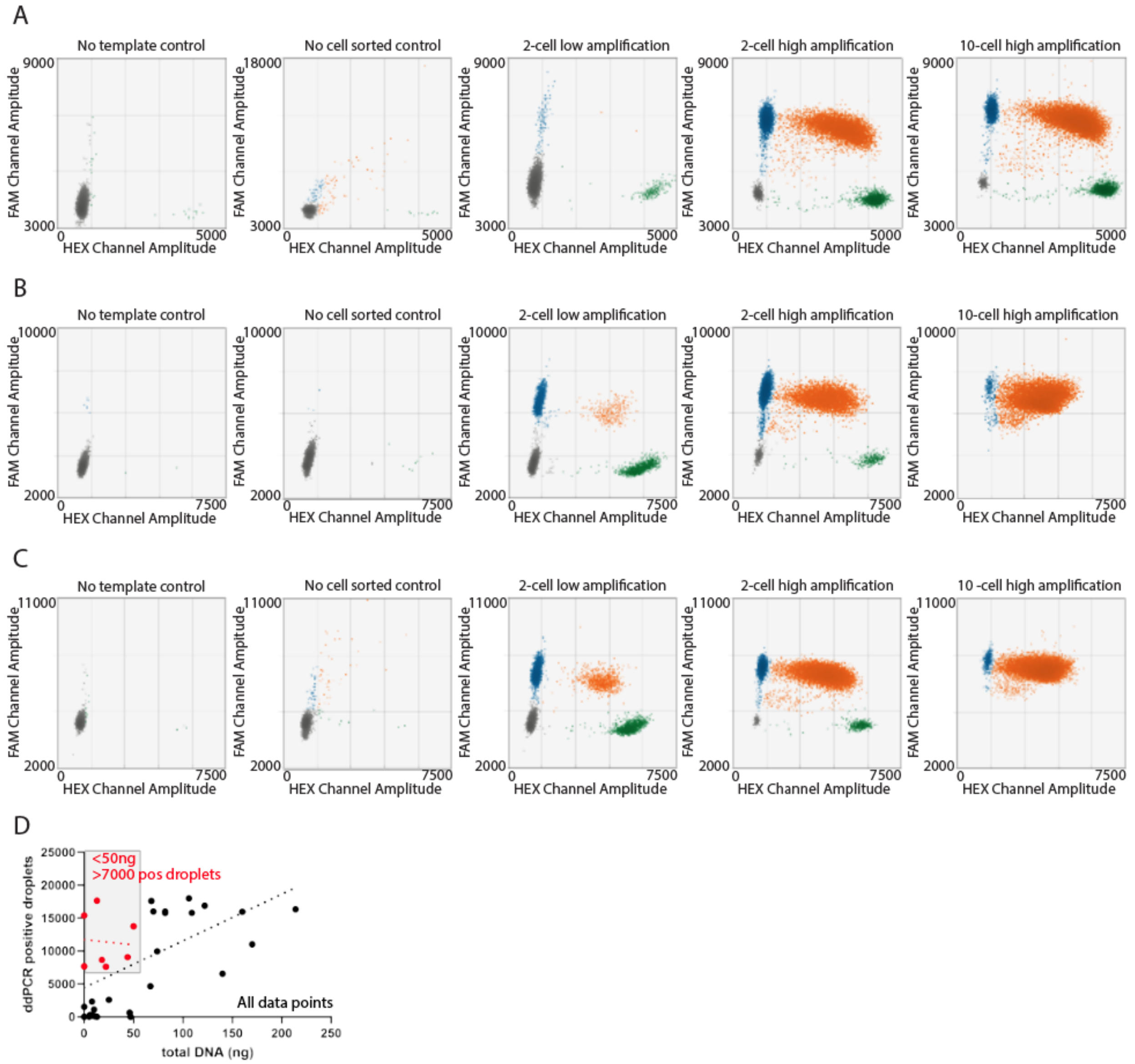


**Figure S7: Parasite-specific ddPCR validates parasite DNA in amplified low-input samples.** Droplet digital PCR (ddPCR) of MALBAC amplified DNA from wells containing 0, 2, and 10 cells. Each colored dot represents a droplet containing a target *P. falciparum* gene (*pfmdr1*: blue, *pfhsp70*: green, both *pfmdr1* and *pfhsp70*: orange, negative population: gray) for a set of representative samples with low and high levels of amplification (listed from left to right). **A**. Untreated *FCR3* samples; no template control, f-a2 (no cell control), f-d3 (2-cell), f-d9 (2 cell), f-d2 (10-cell). **B**. Untreated *Dd2* samples; no template control, u-b6 (no cell control), u-g8 (2-cell), u-d4 (2-cell), u-c2 (10-cell). **C.** DSM1 treated *Dd2* samples; no template control, t-b7 (no cell control), t-f3 (2-cell), t-d4 (2-cell), t-d2 (10-cell). **D**. Comparison of total DNA (from Qubit) and number of positive ddPCR droplet counts from amplified samples where ddPCR was performed. Red points: <50ng but >7000 positive droplets (grey box). Note: all red points are from the treated *Dd2* condition. Linear regression performed in Graphpad PRISM; black dotted line (all data points), p value of 0.0004, $R_2$= 0.34; red dotted line (grey box only), p value of 0.87, $R_2$=0.006.

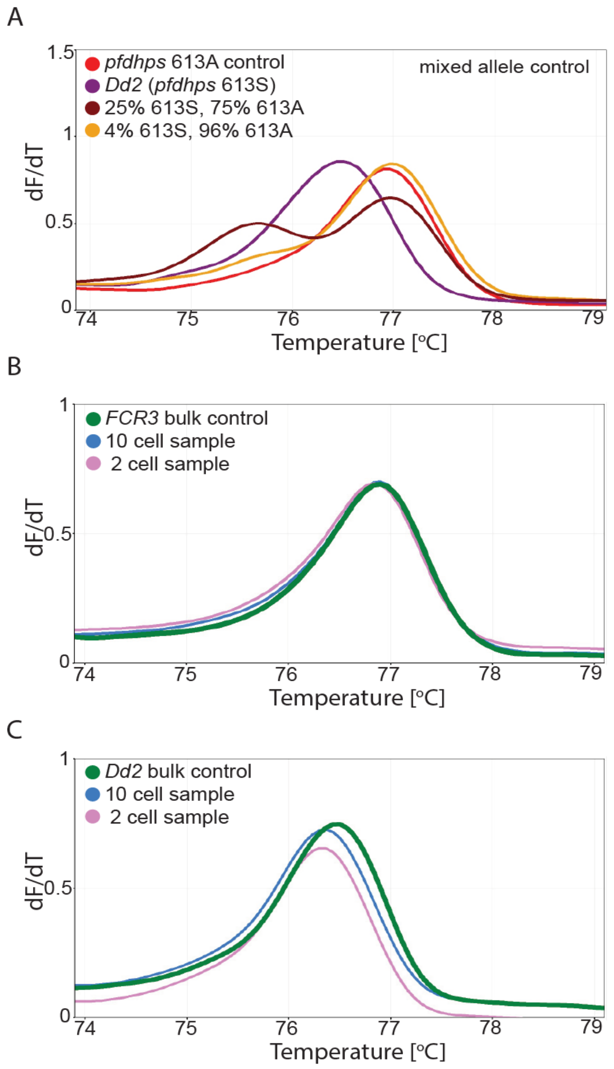


**Figure S8: High-resolution melting assessment displays no evidence of cross-contamination between low-input samples.** PCR followed by high-resolution melting (HRM) analysis of the *P. falciparum pfdhps* gene at position 613 (wildtype control: 613A, mutant allele: 613S) for a set of representative samples. Peaks represent melting temperature for the probe covering the position of interest. dF/dT; change in fluorescence versus temperature. **A**. Comparison of the probe melting temperature for *pfdhps* wt control (613A, *HB3* genomic DNA, same as *FCR3* genotype), mutant genotype (613S, *Dd2* genomic DNA), and mixed genotypes at different proportions to illustrate the impact of mixed alleles on the peak (*HB3* and *Dd2* genomic DNA). **B**. Probe melting temperature for Untreated *FCR3* bulk genomic DNA and MALBAC amplified DNA (f-e7: 2-cell and f-g2: 10-cell, see **Tables S2** and **S3** for sample information) from the Untreated *FRC3* plate. **C**. Probe melting temperature for Untreated *Dd2* bulk genomic DNA and MALBAC amplified DNA (u-e7: 2-cell and u-g2: 10-cell, see **Tables S2** and **S3** for sample information) from the Untreated *Dd2* plate.

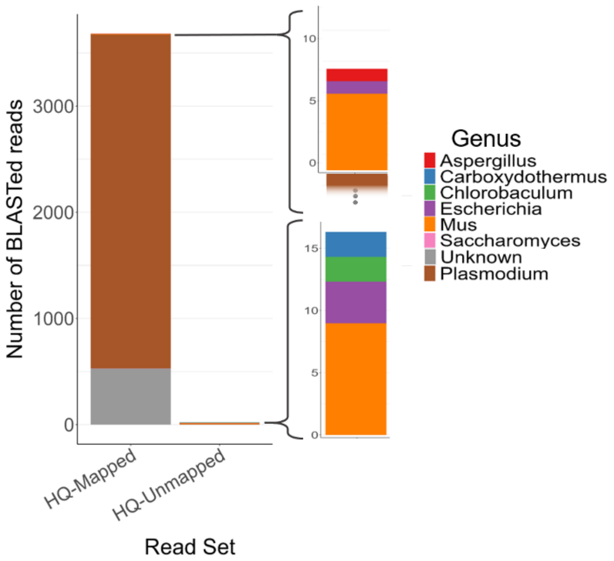


Fig S9. **Comparison of BLAST hits from high quality mapped and unmapped reads from low cell sequencing samples.** Randomly subsampled reads (n = 5000) from either high quality (HQ, >100bp, >30 read q score, <50% N content, see *Materials & Methods*) mapped or unmapped subsets for each low-cell sample. Each stacked bar represents the mean number of reads considered a BLAST hit to the genus per-sample with insets showing proportions unable to be visualized against the total. Unknown genus, reads BLASTed to sequence without a genus tag present in the custom database (see *Materials & Methods*). Mean number of reads for HQ-mapped = 3212 reads, HQ-unmapped = 9 reads, and total unknown genus hits = 911. SD of HQ-mapped *Plasmodium* genus hits = 630 and all other SDs < 12.

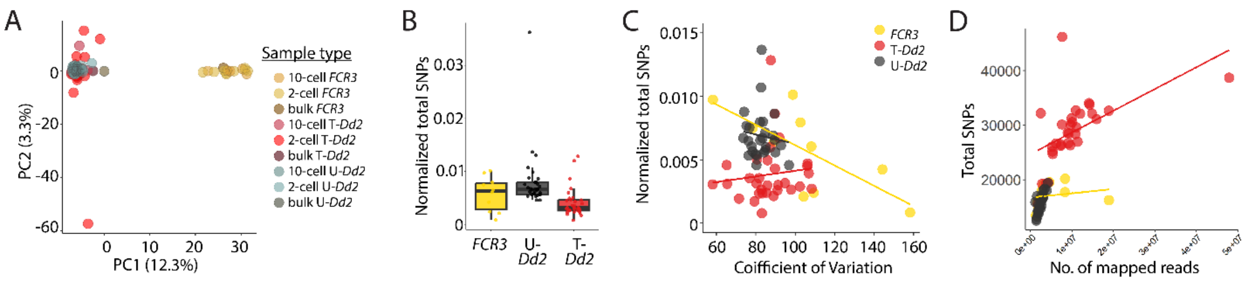

**Figure S10: Single nucleotide polymorphism (SNP) profile of low-input samples shows close relationship with bulk samples and impact of sequencing depth. A**. Principal component analysis of SNP loci from low-input samples (10- and 2-cell) and bulk samples from *FCR3* (yellow) and *Dd2* parental lines (U-*Dd2* (black): untreated *Dd2* samples, T-*Dd2* (red): DSM1-treated samples). All samples are compared to the *Pf*3D7 reference genome (identified a total of 1.1 million SNPs across all samples). Does not include 5 samples excluded due to low coverage (**Table S3**). **B**. Total SNPS for 2-cell samples normalized by total mapped reads. Compared to *FCR3*: NS for untreated *Dd2*, p value of 0.069 for treated *Dd2*; between treated *Dd2* and untreated *Dd2*: p value of 0.000135. **C**. Comparison of total SNPs (normalized to total mapped reads) and coefficient of variation (CV, amplification quality). Untreated *Dd2* $R_2$: 0.01, treated *Dd2* $R_2$: 0.01, *FCR3* $R_2$: -0.51. **D**. Comparison of total SNPs and number of mapped reads, Untreated *Dd2* $R_2$: 0.66, treated *Dd2* $R_2$: 0.27, *FCR3* $R_2$: -0.08.

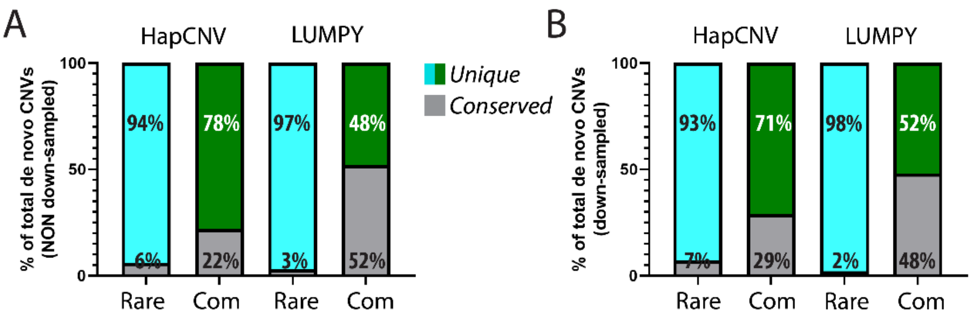


**Figure S11. Rare CNVs across untreated and treated sample types are more often found in unique locations.** Genome location comparison of de novo CNVs across 2-cell untreated and treated samples. Unique: no overlap in location with other de novo CNVs; Conserved: location overlap identified. **A.** Location comparison between untreated and treated samples using all data. **B**. Location comparison between untreated and treated samples using downsampled data (1.3M reads per sample).

A

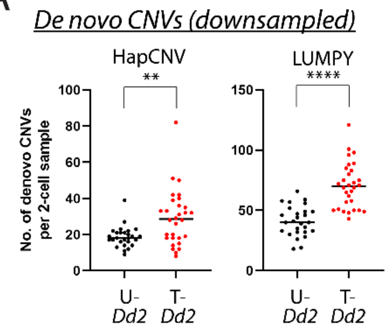


B

| | | Mean CNV count per low-input sample | | | |
|---|---|---|---|---|---|
| **NDS** | CNV type | ***U-Dd2**** | ***T-Dd2**** | Fold dif. | Sig. level |
| LUMPY | de novo | 167.5 | 459.5 | 2.7 | **** |
| | rare | 99.3 | 297 | 3.0 | **** |
| | common | 68.2 | 162.6 | 2.4 | **** |
| | average | nd | nd | 2.7 | - |
| HapCNV | de novo | 16.3 | 30.7 | 1.9 | *** |
| | rare | 1.7 | 7.3 | 4.3 | ** |
| | common | 14.6 | 23.4 | 1.6 | *** |
| | average | nd | nd | 2.6 | - |
| **DS** | CNV type | ***U-Dd2**** | ***T-Dd2**** | Fold dif. | Sig. level |
| LUMPY | de novo | 52.3 | 77.6 | 1.5 | **** |
| | rare | 33 | 54.9 | 1.7 | **** |
| | common | 19.3 | 22.6 | 1.2 | * |
| | average | nd | nd | 1.4 | - |
| HapCNV | de novo | 18.7 | 29.1 | 1.6 | ** |
| | rare | 3.3 | 12.2 | 3.7 | *** |
| | common | 15.4 | 16.9 | 1.1 | ns |
| | average | nd | nd | 2.1 | - |

C

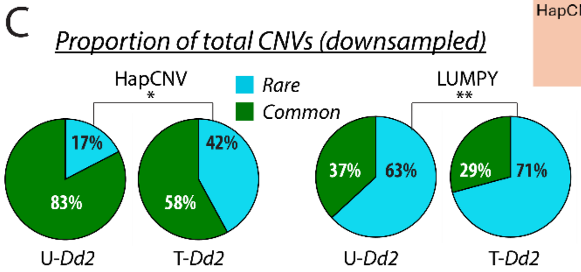


**Figure S12. Downsampled data shows consistent increase in CNVs following replication stress. A.** Detection of de novo CNVs (common and rare combined) from untreated (U-*Dd2*) and treated (T-*Dd2*) 2-cell samples using two CNV analysis methods on downsampled reads (unpaired T-test with two tailed Welch's correction, p values: 0.0014 for HAPCNV and <0.0001 for LUMPY). Line at mean value for each dataset. **B**. Summary of results from all data (non-downsampled, NDS, blue) and downsampled data (DS, orange) from 2-cell samples in untreated (U-*Dd2*) and treated (T-*Dd2*) samples. *, mean calculated by Graphpad PRISM during Welch's test; nd, not determine. Fold difference is calculated as treated/untreated CNV counts. The de novo category represents the sum of rare and common CNVs. Significance levels assess difference: ****, p value < 0.0001; ***, <0.001; **, <0.01; *, <0.1; ns, not significant; -, not applicable. **C**. Proportion of total CNVs detected as rare and common from downsampled reads (1.3M total reads); pie charts plot the mean but statistics are calculated using all data points from the rare CNV category (p values: 0.02 for HAPCNV and 0.004 for LUMPY).

**Table S1.** *Pf*MALBAC version 2 improves coverage of intergenic regions and decreases amplification bias.

| Conditions* | Pre-amplification | | Coverage (X) | GC % | Coverage breadth | | | |
|---|---|---|---|---|---|---|---|---|
| | Polymerase | Random primer[$] | | | Whole genome | Genic regions | Intergenic regions | Coefficient of Variation (CV^) |
| *Pf*MALBAC_v1 | *Bst* | v1 | 1.6 | 24.7 | 26.5% | 41.7% | 4.0% | 126 |
| *Pf*MALBAC_v2 | *Bst* | v2 | 1.7 | 23.1 | 29.3% | 45.4% | 5.2% | 109 |
| | *Bst* | v2 | 1.7 | 22.6 | 22.2% | 32.8% | 6.6% | 131 |
| | *Bsu* | v1 | 1.7 | 23.0 | 34.0% | 50.2% | 9.9% | 65 |
| | *Bsu* | v2 | 1.7 | 22.7 | 35.3% | 52.0% | 10.5% | 78 |
| | *Bsu* | v2 | 1.7 | 22.4 | 35.8% | 52.4% | 11.2% | 69 |

*All amplification conditions were performed on *Dd2* ring-stage *P. falciparum* parasites. *Bst* amplification is performed at 64°C and *Bsu* amplification is at 37°C.
^CV is the coefficient of variation of normalized read abundance as in Liu et al. (42).
[$]Random primer version 1 (v1): 5'GTGAGTGATGGTTGAGGTAGTGTGGAGNNNNNTTT 3'; version 2 (v2): 5'GTGAGTGATGGTTGAGGTAGTGTGGAGNNNNNNNNNNTTT 3'

**Table S2**: Details of library preparation for low-input sequencing. **IN SEPARATE FILE**

**Table S3**: Illumina short-read sequencing statistics for low-input samples. **IN SEPARATE FILE**

**Table S4.** CNV detection in low-input samples from total reads using two methods. **IN SEPARATE FILE**

**Table S5**. Summary of genes from high confidence CNV regions. **IN SEPARATE FILE**

**Table S6.** Properties of high-confidence CNV breakpoints. **IN SEPARATE FILE**

**Table S7**. Lack of protein class enrichment from high confidence CNV regions. **IN SEPARATE FILE**
